# Phenotypic Plasticity of Symbiotic Organ Highlight Deep-sea Mussel as Model Species in Monitoring Exploitation of Deep-sea Methane Hydrate

**DOI:** 10.1101/2022.08.11.503589

**Authors:** Mengna Li, Hao Chen, Minxiao Wang, Zhaoshan Zhong, Chao Lian, Li Zhou, Huan Zhang, Hao Wang, Lei Cao, Chaolun Li

## Abstract

The rapid progress of deep-sea methane hydrate exploration and exploitation calls for a thorough evaluation of its potential impact on local chemosynthetic ecosystems, particularly on endemic species. As one dominant megafauna in cold seeps, the deep-sea mussels mainly rely on methanotrophic endosymbionts for nutrition and therefore could serve as a promising model in monitoring the exploitation of methane hydrate. However, knowledge on the long-term responses of deep-sea mussels to environmental stresses induced by methane hydrate exploitation, especially to methane reduction and deprivation, is still lacking. Here, we set up a laboratory system and cultivated methanotrophic deep-sea mussel *Gigantidas platifrons* without methane supply to survey the phenotypic changes after methane deprivation. While the mussels managed to survive for more than 10 months after the methane deprivation, drastic changes in the metabolism, function, and development of gill tissue, and in the association with methanotrophic symbionts were observed. In detail, the mussel digested all methanotrophic endosymbionts shortly after methane deprivation for nutrition and remodeled the global metabolism of gill to conserve energy. As the methane deprivation continued, the mussel replaced its bacteriocytes with ciliated cells to support filter-feeding, which is an atavistic trait in non-symbiotic mussels. During the long-term methane deprivation assay, the mussel also retained the generation of new cells to support the phenotypic changes of gill and even promoted the activity after being transplanted back to deep-sea, showing the potential resilience after long-term methane deprivation. Evidences further highlighted the participation of symbiont sterol metabolism in regulating these processes, which might be one direct cue for the mussels to respond to methane deprivation. These results collectively show the phenotypic plasticity of deep-sea mussels and their dynamic responses to methane deprivation, providing essential information in assessing the long-term influence of methane hydrate exploitation.

## 1. Introduction

Methane hydrate (also known as natural gas hydrate or combustible ice, hereinafter referred to as the hydrate) is a new unconventional alternative energy source due to the wide distribution, abundant reserves, and high fuel efficiency (Boswell, 2009). It is estimated that the global reserve of methane hydrates is up to 3,000 trillion cubic meters (a previous estimation of 20,000 trillion cubic meters), which are far more than the global reserves of conventional fossil fuel resources as well as the shale gas resources (Chong et al., 2016; Milkov, 2004). On the meanwhile, the methane hydrates contain more carbon than all other fossil fuels combined, making it potential alternative of traditional fossil fuels to mitigate global warming induced by CO_2_ emissions (Chong et al., 2016; Makogon et al., 2007). Therefore, although 97%-99% of the hydrates are located in the ocean, demand for the exploration and exploitation of methane hydrate is increasing. To date, many countries have carried out their own explorations and trial mining of the hydrate in the deep ocean (Liu et al., 2019b). However, the exploration and exploitation of methane hydrate are still facing multiple challenges, including technological challenges, stability and safety, cost and economics, regulatory and legal frameworks (Chong et al., 2016; Liu et al., 2019a). In addition to these issues, environmental impacts during the mining of methane hydrates also need to be thoroughly assessed as the methane may escape during drilling and reach to the atmosphere (Zhao et al., 2017). Besides, whether the mining of methane hydrates could result in submarine slope failure has drawn much attention as well since the hydrates are usually buried in the sands or muds beneath the seafloor (Kwon and Cho, 2012).

It is important to note that chemosynthetic communities are frequently found alongside methane hydrates in the deep sea, particularly when the hydrates form substantial and solid mounds directly on the seafloor or are buried in near-surface sediments (Feng et al., 2018; Fisher et al., 2000; Sen et al., 2018; Zhang et al., 2023). The chemosynthetic communities, since their first discovery in 1980s, have been observed to sustain a wide range of unique and native marine megafauna, such as tubeworms, mussels, clams, and shrimps, through close symbiotic associations (Sogin et al., 2020). It is therefore necessary to assess the impact of methane hydrate mining to these communities, especially the benthic megafauna. To date, many efforts with *in-situ* experiments and long-term observations have been made to investigate the influence of deep-sea mining on the biodiversity of microbial or megafauna communities, providing substantial knowledge on the ecological impact of light, noise, sedimentation and other stresses caused by the mining (Borowski and Thiel, 1998; Ingole et al., 2005; Spearman et al., 2020; Vonnahme et al.; Washburn et al., 2023). Among the potential environmental changes, the reduction and depletion of methane hydrates is undoubtedly the most significant one as the oxidation of methane provides primary sources of energy and carbon for both the seep ecosystem. However, the long-term effects of the reduction and depletion of methane hydrates on the biodiversity of local community and on the physiology of chemosynthetic bacteria and megafauna are still insufficiently investigated due to the technical challenges on the *in-situ* experiments and observations.

As with many other endemic species in cold seep ecosystems, deep-sea Bathymodiolinae mussels are known to harbor methanotrophs and/or thiotrophs in their gill tissue as their major source of nutrition (Dubilier et al., 2008). The deep-sea mussels are estimated to be diverged from their non-symbiotic shallow-water relatives 110 million years ago and have acquired specialized cells (known as bacteriocytes) in the gill tissue to harbor the chemosynthetic endosymbionts during the migration to deep-sea (Kiel and Amano, 2015; Miyazaki et al., 2010). The bacteriocytes constitute more than 20% of gill epithelial cells (in adult mussels) and can acquire the symbionts horizontally from local environments or from adjacent bacteriocytes (Franke et al., 2021; Wang et al., 2023; Wentrup et al., 2014). While the Bathymodiolinae mussels rely mostly on their endosymbionts for nutrition, they can also acquire organic matter through filter-feeding (Martins et al., 2008; Page et al., 1990), and may even lose their symbiotic association if their chemosynthetic sources are reduced while organic matter is still accessible (Rodrigues et al., 2015). In addition, deep-sea mussels can also adjust the types or relative abundances of endosymbionts based on the local environment, and can even regain their symbiotic association after the complete decolonization of symbionts when the methane/sulfide supply recovers (Halary et al., 2008; Kádár et al., 2005; Rodrigues et al., 2015; cker et al., 2021). The diversity and plasticity of symbiotic associations in deep-sea mussels have made them promising model in monitoring the potential influences, and therefore determining threshold sensitivities, of methane hydrate mining to megafauna. Recent studies further showed that majority genes involved in methane metabolism of methanotrophs were repressed during methane deprivation, which further result in the arrest of growth (Tavormina et al., 2017). For the mussel hosts, a promoted lysosome-mediated digestion of methanotrophic symbionts could be found in the bacteriocytes after the methane deprivation, following the activation of immune and apoptotic processes of gill tissue (Détrée et al., 2019; Sun et al., 2022; Wang et al., 2019). However, knowledge on the long-term influence of deep-sea mussels during methane reduction and deprivation is still lacking due to the short period of time examined in previous studies, which typically ranges from one week to one month (Sun et al., 2022; Wang et al., 2019). Since the deep-sea mussels are reported to survive for years even after their complete loss of symbionts, we hypothesize that deep-sea mussels might be able to adjust the function and development of gill tissue dynamically in response to methane deprivation and therefore sustain the long survival. To validate our hypothesis and clarify the physiological changes of deep-sea mussels in simulated methane hydrate mining, we conducted a 10-month methane deprivation assay under laboratory conditions with methanotrophic deep-sea mussel (*Gigantidas platifrons*) and then assessed the phenotypic changes of gill tissue using proteomics and metabolomics data and EM analysis. We also conducted a transplantation assay of decolonized mussels back to deep-sea to assess the potential resilience of mussel by examining the generation of gill cells. It is our hope that our results will deepen our understanding of the dynamic response of deep-sea mussels to long-term methane deprivation and provide useful information in evaluating the potential influences and threshold sensitivities of methane hydrate mining.

## 2. Materials and methods

### 2.1. Mussel collection and methane-deprivation assay

The sample information and experimental design of this study are summarized in Fig. 1A. In detail, *G. platifrons* mussels were collected from a seepage region of a cold seep (22°06′N, 119°17′E; 1,113 m depth) in the South China Sea using the “Faxian” remotely operated vehicle, operated on the R/V “Kexue” during a cruise in 2017. The mussels were transferred to the deck for less than 1 h using an isobaric and/or isothermal biobox that could shelter them from depressurization (except for the isothermal biobox) and elevated temperature during recovery (Chen et al., 2021). Once aboard, the gill tissues from six mussels were collected to indicate the *in-situ* state (the InS group, collected using an isobaric isothermal biobox). The remaining mussels (collected using an isothermal biobox) were transferred to the onboard aquarium (which had continuously recirculating seawater and was under atmospheric pressure) for methane deprivation.

**Figure 1.**
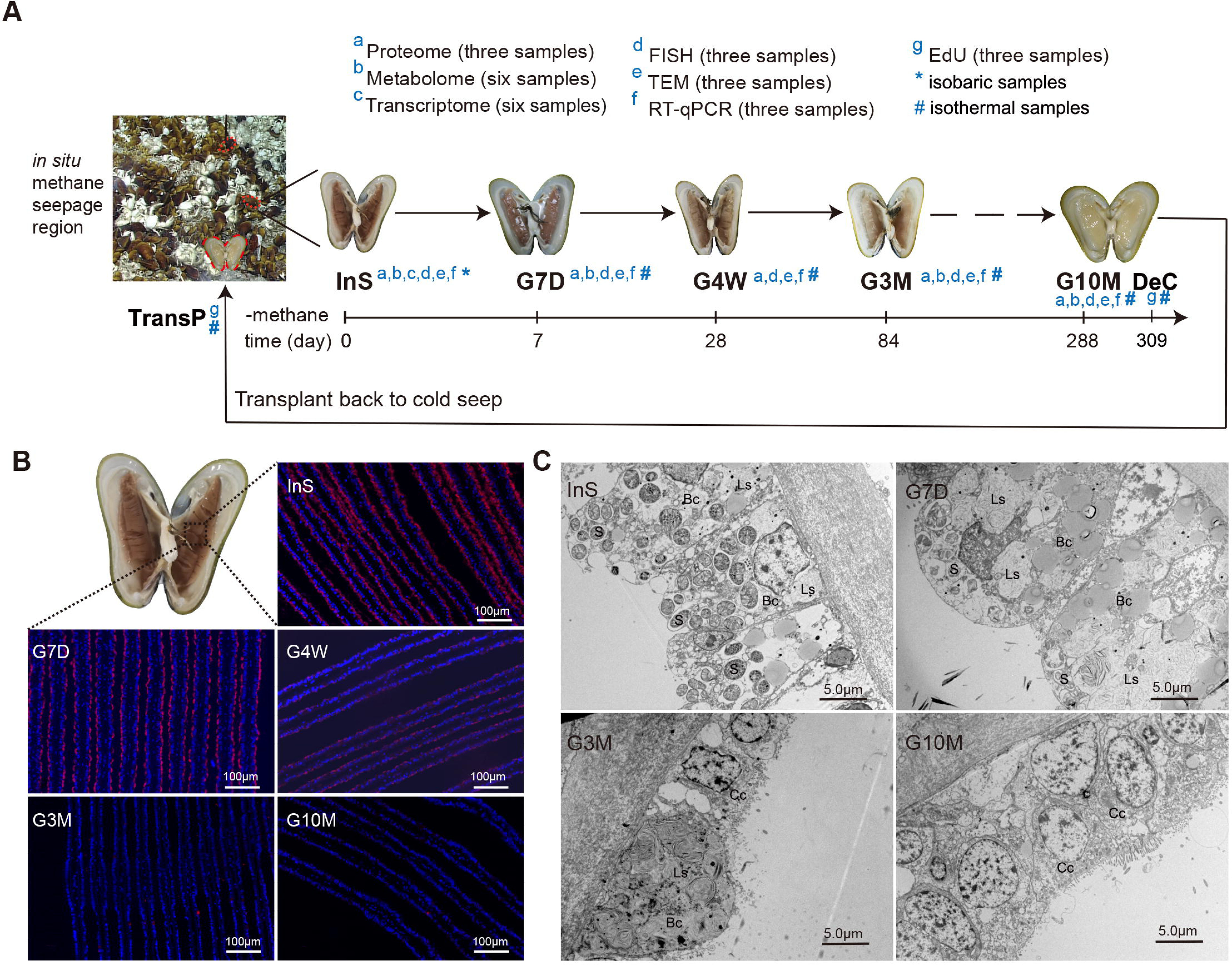
Phenotypic changes of gill tissue under methane deprivation. **(A)** A schematic drawing of the experimental workflow. To assess phenotypic changes of gill tissue during methane deprivation, methanotrophic deep-sea mussels were acclimated in a laboratory circulating system without methane supply for different periods of time. Mussels on the first day (InS), at 7 days (G7D), at 4 weeks (G4W), at 3 months (G3M), and at 10 months (G10M) of acclimation were sampled for subsequent analysis. To assess developmental plasticity of gill tissue, decolonized mussels cultivated onboard for 309 days (DeC group) were transplanted back to cold seeps after EdU labelling and retrieved five days (TransP group) later for further analysis. **(B)** Fluorescence *in situ* hybridization (FISH) assay of mussel gills. The Cy3-labelled symbiont-specific probe (designed based on the pmoB gene) was employed to indicate endosymbiotic methanotrophs (in red), while nuclei were stained using 4′, 6-diamidino-2-phenylindole (DAPI, in blue). A gradual termination of Cy3 signals was observed as the methane deprivation assay continued, indicating the depletion of symbionts. **(C)** Transmission electron microscope (TEM) analysis of cross-sectioned gill tissues. A marked decline in the symbiont population (S) and increases of lysosomes (Ls) in bacteriocytes (Bc) were observed shortly after the methane deprivation assay (G7D group). Bacteriocytes (Bc) were then gradually replaced by ciliated cells (Cc) in the mussels of of the G3M and G10M groups.

During onboard methane deprivation, surface seawater above the cold seep was collected and adjusted to 3.5 ± 0.5°C and 34.5 ± 0.5‰ salinity to be consistent with the *in situ* environment (Cao et al., 2021). The seawater inside the aquarium continuously recirculated between a refrigerator, a sand filter tank, and a protein skimmer, and half of it was replaced every day. After 1 month in the onboard facility, the mussels were transferred into a recirculating system inside the laboratory with same environmental parameters and recirculating schedule (using seawater pumped from Jiaozhou Bay, filtered with sand and sterilized using ultraviolet irradiation). On the 7^th^, 28^th^, 84^th^, and 288^th^ days post acclimation, a total of 24 mussels were randomly collected and subjected to proteomic, metabolic, and morphological analysis (designated as the G7D, G4W, G3M, and G10M groups, respectively, with six replicates for each group). In addition, since the main goal of our study is to characterize the physiological response of mussels to methane deprivation and considering that it remains a challenge to cultivate deep-sea mussels under pressurized condition and the depressurization could alter the global metabolism and gene expression of endosymbionts, we treated the mussel holobionts with a final concentration of 50 mg/L of ampicillin and kanamycin during the first month after recovery to suppress and silence any biased expression of endosymbionts. The treatment had no influence on the symbiont abundance as well as the mussel expression according to results of an *in situ* antibiotics treatment assay (see Supplementary text). For all sampled mussels, a portion of the gill filaments was fixed with 4% paraformaldehyde or a mixture of paraformaldehyde and glutaraldehyde (2%/2.5%) for the subsequent fluorescence *in situ* hybridization (FISH) and transmission electron microscopy (TEM) assays. The remaining gills were frozen with liquid nitrogen and stored at −80°C for symbiont quantification, proteome analysis (only gills from three mussels each group were used), and metabolome analysis (the G4W group was excluded from the analysis).

### 2.2. In situ transplantation assay and onboard pressurization assay

To confirm the potential resilience of mussel gills, an *in situ* transplantation assay (with EdU-labeling) was performed by returning decolonized mussels to cold seeps. In detail, on the 309^th^ day after mussel recovery, a total of 18 live mussels that had been cultivated continuously in the laboratory recirculating system (in the same aquarium and subjected to the same methane deprivation) at were injected with 5-ethynyl 2′-deoxyuridine (EdU, 5×10^−9^ mol/individual) in the adductor muscle to label the newly generated gill cells. Of the injected mussels, six were added back into and continuously cultured in the recirculating system (the DeC group, an abbreviation of decolonized mussels), while the remaining mussels were returned to the methane seepage region in the cold seep using the isothermal biobox (the TransP group, an abbreviation of transplantation back to seep). The EdU-treated mussels were collected 5 days later for gill tissue sampling and fixed with paraformaldehyde or paraformaldehyde/glutaraldehyde for EdU staining and TEM analysis.

To assess the potential influences of antibiotics on mussel holobionts, we also conducted antibiotic treatment with the same dosage with an onboard experiment (50 mg/L ampicillin and kanamycin) for 7 days *in situ* at the cold seep. A multipurpose *in situ* sampling device was used to shelter the samples from depressurization and heat, and six antibiotics-treated mussel samples (InS_At group) were fixed with RNAsafer stabilizer reagent (Omega Bio-Tek, Norcross, GA, USA) before their retrieval back to the deck. As the control group, another six mussels (InS group) from the seepage region were also collected and treated with RNAsafer *in situ*. For mussels in these two groups, only meta-transcriptome analysis was conducted.

To assess the potential influences of depressurization on the symbiont abundance, a re-pressurization assay was also conducted using pressurized cabins. In short, a total of 12 deep-sea mussels collected in an isothermal way were acclimated for up to six days with an addition of 2 MPa gas mixtures (30% methane, 50% nitrogen, and 20% air) in filtered seawater and sampled at 24 h, 72 h, and 144 h post treatment (2 MPa Group). As controls, another 16 mussels were acclimated under atmospheric pressure conditions (supplemented with the same gas mixtures) and sampled simultaneously with the 2 MPa group. All samples were dissected for gill tissue and were subjected to genome DNA extraction and qRT-PCR of *pmoA* genes.

### 2.3. Proteome analysis

Proteome analysis was conducted with assistance by PTM Biolabs (Hangzhou, China). The proteome analysis methods are described in detail in the supplementary material (see supplementary text). In brief, gill proteins from three individuals from each group were extracted and digested by trypsin. The peptides obtained by trypsin digestion were subsequently labeled as directed by the tandem mass tag (TMT) kit (PTM Biolabs). The labeling peptides were then fractionated by high pH reverse-phase high-performance liquid chromatography and further subjected to liquid chromatography with tandem mass spectrometry (LC-MS/MS) analysis. LC-MS/MS spectra were searched against the *G. platifrons* genome data concatenated with the reverse decoy database. For the identification of differentially expressed proteins, the expression values across all samples were first quantified and the mean values of each group were used for subsequent comparison. Proteins were considered to be differentially expressed only when the fold changes between two given groups were greater than 1.2-fold, with a p value less than 0.05 (two-tailed *t*-test). Gene Ontology (GO) and KEGG annotations of the *G. platifrons* proteome were derived from the integrated analysis by non-redundant protein sequences of the NCBI, the KEGG database, UniProt, Pfam, and InterProScan as described in the supplementary materials.

### 2.4. Metabolome analysis

The detailed metabolome analysis methods are described in the supplementary material. In brief, gill tissues from six mussels in each group were treated with 80% methanol to release all metabolites. After centrifugation at 12,000 g and 10°C for 15 min, the supernatants were divided into four parts and subjected to liquid chromatography–mass spectrometry (LC-MS), gas chromatography/mass spectrometry (GC–MS), and the corresponding quality control analysis.

The raw spectra obtained by LC-MS and GC-MS were decoded by MetaboAnalyst (https://www.metaboanalyst.ca/MetaboAnalyst/ModuleView.xhtml), and all identified metabolites were annotated with PubChem, and the KEGG database. For the identification of differential metabolites, a PCA analysis was first conducted to validate the consistency of parallel samples within the same group using the OmicStudio tools (https://www.omicstudio.cn/). Then, the differentially abundant metabolites were screened based on the variable importance for the projection (VIP) of OPLS-DA test. Metabolites were considered to be differentially abundant when VIP > 1 and *p*-value < 0.05 (two tailed *t*-test).

### 2.5. Meta-transcriptome analysis

Meta-transcriptome analysis was performed by Genedenovo (Guangzhou, China) as described previously (Chen et al., 2021). In brief, gill tissues from six InS mussels and six *in situ* antibiotic-treated mussels were subjected to RNA extraction using TRIzol reagent (Invitrogen, MA, USA), and RNA quantification, mRNA purification, cDNA synthesis, adaptor ligation, cDNA library construction, and sequencing using the Illumina HiSeq 2500 platform with paired-end reads were conducted by the Gene Denovo Biotechnology Co (Guangzhou, China). Differentially expressed genes between the two groups were screened using the edgeR package in the R software when the fold change > 2.0 and false discovery rate < 0.05. KEGG annotation and enrichment analysis were likewise conducted as described above for the proteome analysis.

### 2.6. Weighted gene co-expression network analysis

Weighted gene co-expression network analysis (WGCNA) was conducted to identify proteins with correlated expression patterns in all groups. Co-expression networks were constructed using the WGCNA package (v1.47) in the R software after incorporating the protein abundance (power = 6). The correlation coefficients of samples were then calculated based on the module eigengenes. The intramodular connectivity and module correlation degree of each gene was also calculated by the R package of WGCNA to determine the hub genes. Finally, the networks were visualized using Cytoscape (v3.8.2).

### 2.7. Quantitative real-time PCR assay for symbiont quantification

As a classic way to detect relative expression of target genes, quantitative real-time PCR (qRT-PCR) is also widely used to assess the relative changes in symbiont abundance (Sun et al., 2017; Wang et al., 2019; Yu et al., 2019). Here, the genomic DNA of gill tissues was first extracted using a mollusk DNA extraction kit (Omega Bio-tek, Norcross, GA, USA) for the qRT-PCR template. The methane monooxygenase subunit A gene (*pmoA*), a marker gene in methanotrophic symbionts, was used to indicate the symbiont abundance, while the mussel actin gene was used as an internal control. The primers and qRT-PCR programs used were the same as those described in previous reports (Chen et al., 2019). For statistics analysis, six biological replicates were employed for each group, and the relative abundance was calculated using the 2^−ΔΔCt^ method and given as the mean ± s.d. A Kruskal–Wallis test followed by Dunn’s multiple comparison was employed to determine the significant differences between groups.

### 2.8. Transmission electron microscopy analysis

TEM analysis of the gills was conducted following a method described in a previous study (Wang et al., 2021). In brief, gill samples fixed by paraformaldehyde-glutaraldehyde were treated with 1.0% osmium tetroxide (OsO_4_) for staining and then dehydrated with alcohol. After infiltration with acrylic resin, the samples were finally embedded and sectioned with a thickness of 70 nm using an ultramicrotome (EM UC7, Leica, Vienna, Austria). After staining by uranyl acetate and lead citrate, the gill cell ultrastructure was imaged with a transmission electron microscope (HT7700, Hitachi, Tokyo, Japan).

### 2.9. Fluorescent in situ hybridization assay and EdU staining assay

The fluorescent *in situ* hybridization (FISH) assay of symbionts was conducted using gill filament samples from the InS, G7D, G3M, and G10M groups according to the method described previously (Halary et al., 2008; Wang et al., 2021). In brief, gill samples fixed by paraformaldehyde were first dehydrated and embedded in paraffin. After they were cross-sectioned to 7 μm in thickness, sections were rehydrated in xylene twice for 10 min, and then in a 1:1 mixture of 100% ethanol and xylene, 100% ethanol (twice), 95% ethanol, 85% ethanol, and finally 75% ethanol, for 5 min each.

After being permeabilized with 1×PBST (1×PBS with 0.1% Tween 20) for 15 min, the gill tissues were hybridized with 0.5 ng/μL Cy3-labeled *pmoB* gene probe (5’CGAGATATTATCCTCGCCTG3’), a methanotroph specific probe, in hybridization buffer (0.9 M NaCl, 0.02 M Tris-HCl, 0.01% sodium dodecyl sulphate, and 30% formamide) for 1 h at 46℃, followed by washing with washing buffer (0.1 M NaCl, 0.02 M Tris HCl, 0.01% SDS and 5 mM EDTA) for 15 min at 48℃. After staining the nucleic acids with 4′, 6-diamidino-2-phenylindole (DAPI), gill sections were imaged with a fluorescence microscope (Nikon ECLIPSE Ni, Tokyo, Japan) to demonstrate the spatial distribution of symbiotic methanotrophs. EdU staining was also conducted with the paraffin sections. In brief, after treatment with 4% paraformaldehyde and dehydration with ethanol, EdU-labeled gills were embedded in paraffin and sectioned using a microtome (Leica, Heidelberg, Germany) to a thickness of 7 μm. The EdU staining was conducted using the Click-iT Plus EdU Imaging Kit (Thermo Fisher, Waltham, MA, USA) and visualized with a fluorescence microscope (Nikon ECLIPSE Ni, Tokyo, Japan) after deparaffinization and rehydration.

## 3. Results

### 3.1. Phenotypic changes of gill tissue during the laboratory methane deprivation

To simulate methane hydrates mining and assess the long-term phenotypic changes of the deep-sea mussel during methane reduction and deprivation, a laboratory recirculating aquarium system with an environment similar to that of a cold seep (including similar temperatures and salinities, but without a methane supply) was set up and employed in the present study (Fig. 1A). Using this system, mussels cultivated for different periods of methane deprivation were successfully collected, including mussels cultivated for 7 days (G7D), 4 weeks (G4W), 3 months (G3M), and 10 months (G10M). Although the mussels managed to survive more than 10 months after methane deprivation, drastic changes in the morphology of gill tissue and in the symbiotic association with methanotrophs were observed. In detail, the gill tissue gradually became thin and bleached during the laboratory treatment. Quantitative real-time PCR (qRT-PCR), FISH, and TEM analyses further showed that both the density and overall number of symbionts decreased continuously under laboratory methane deprivation, and were almost no longer able to be detected after the third month of laboratory cultivation (including the G3M and G10M groups, Fig. 1B, C, Fig. S2A). Accompanied by the decolonization of symbionts, it was also noticeable that the relative number and volume of secondary lysosomes in bacteriocytes, as revealed by TEM analysis, increased drastically in decolonized mussels of the G7D and G4W groups in comparison with control mussels (InS group) (Fig. 1C). As methane deprivation and symbiont losses continued, the bacteriocytes disappeared gradually in the gill tissue after the third month (including the G3M and G10M groups) while the voids left along the gill filaments were occupied by ciliated cells (Fig. 1C).

Proteome and metabolome of the symbiotic (InS group) and decolonized mussels (G7D/G4W, G3M, and G10M) were then conducted to clarify the molecular responses of deep-sea mussel along with these physiological changes during the methane deprivation. Consequently, a total of 7821 host proteins occupying 22.58% of total encoding genes in *G. platifrons*, and 878 endosymbiont proteins occupying 14.21% of their total encoding genes were identified from the proteome data of all groups (Table S2, Supplementary text). Based on the aims and goals of the present study, only quantifiable host proteins (a total of 6611 proteins) were subjected to subsequent analysis. Principal component analysis (PCA) and Pearson correlation analysis demonstrated good consistency among samples of the same group, verifying the repeatability and reliability of the proteomic data (Fig. S2C, D). In addition, by screening the differentially expressed proteins of each group in comparison with the InS group, it was found that the expression profiles of the majority decolonized groups (the G7D, G4W, and G10M groups, with the exception of the G3M group) were distinctly different from those of symbiotic mussels (InS group), while the G7D and G4W groups were highly similar to each other, sharing the majority of differentially expressed proteins (Fig. S2E). The distinct expression profiles demonstrated the vigorous response of deep-sea mussels during methane deprivation and indicated that the hosts might adopt different strategies at the early-stage (the G7D and G4W groups), middle-stage (the G3M group), and long-term (the G10M group) of methane deprivation-induced symbiont decolonization. Consistent with the above findings, functional enrichment analysis of the differentially expressed proteins also showed that the hosts employed distinct biological processes at different stage of methane deprivation, ranging from cell metabolism to cytoskeleton/organelle rearrangement and cell proliferation/differentiation (Supplementary text, Fig. S3, Fig. S4, and Table S3). For example, lysosome and peroxisome-mediated biodegradation and pyruvate metabolism were enriched in up-regulated proteins in the G7D and G4W groups in comparison with InS group, while the bacterial invasion of epithelial cells, endocytosis, regulation of cytoskeleton organization, and cholesterol metabolism were enriched in the down-regulated proteins (Fig. S3AB, Fig. S4AB). The lysosome-mediated biodegradation was enriched in the down-regulated proteins of the G3M group in comparison with the InS group, while processes such as protein digestion and absorption, central carbon metabolism, amino acid transportation, and cell differentiation were enriched in the up-regulated proteins (Fig. S3C, Fig. S4C). At the 10^th^ month after methane deprivation (the G10M group, in comparison with the InS group), processes such as mitophagy, amino acid biodegradation, glycolysis/gluconeogenesis, pyruvate metabolism, the TCA cycle, and oxidative phosphorylation were enriched in the up-regulated proteins, while processes such as endocytosis, protein digestion and absorption and DNA replication were enriched in the down-regulated proteins (Fig. S3D, Fig. S4D). In consistency with the proteomic data, metabolome results also showed distinct metabolic patterns between groups. Noticeably, metabolites involved in glycolysis/gluconeogenesis, pyruvate metabolism, the TCA cycle, protein digestion and absorption, and lipid turnover were found changing significantly during the methane deprivation (Fig. S5, Table S4, and Table S5). Among these processes, the dynamic modulations of the cell metabolism, cytoskeleton/organelle rearrangement, and cell proliferation/differentiation were consistent with the morphological changes observed in the FISH and TEM assays.

### 3.2. Lysosome-mediated symbiont digestion highlighted metabolic adjustment of gill at the early stage of methane deprivation

Among the morphological changes of bacteriocytes that occurred shortly after methane deprivation, the depletion of endosymbionts was of great importance as they served as the major nutrition sources of the hosts. With high-resolution TEM images of bacteriocytes, more methanotroph debris was observed in the lysosome, indicating the participation of lysosome-mediated digestion in the symbiont depletion (Fig. 2A). In support of this speculation, the lysosome pathway was significantly up-regulated in the G7D group compared to the InS group (Fig. S3A). In detail, it was noted that the majority (11/13) of the differentially expressed proteins involved in the lysosome were up-regulated in the G7D group (Fig. 2B). Accompanied by the enhancement of lysosome activity, several key enzymes (15 out of 20) involved in the β-oxidation of fatty acids of the peroxisome, including acyl-CoA oxidase (ACOX1), enoyl-CoA hydratase 2 (ECH2), carnitine O-palmitoyltransferase 1 (CPT1A), and (3R)-3-hydroxyacyl-CoA dehydrogenase (HSD17B4), were also found to be promoted (Fig. 2C), resulting in the accumulation of some intermediates of fatty acid oxidation such as lysophosphatidylcholine (LPC), lysophosphatidylethanolamine (LPE), and carnitine (Fig. 2D, Table S5). These results collectively confirmed the participation of lysosome-mediated digestion in methane deprivation-induced symbiont depletion, which is an essential way for the host to obtain nutrition when symbionts are no longer of use to the host due to methane deprivation.

**Figure 2.**
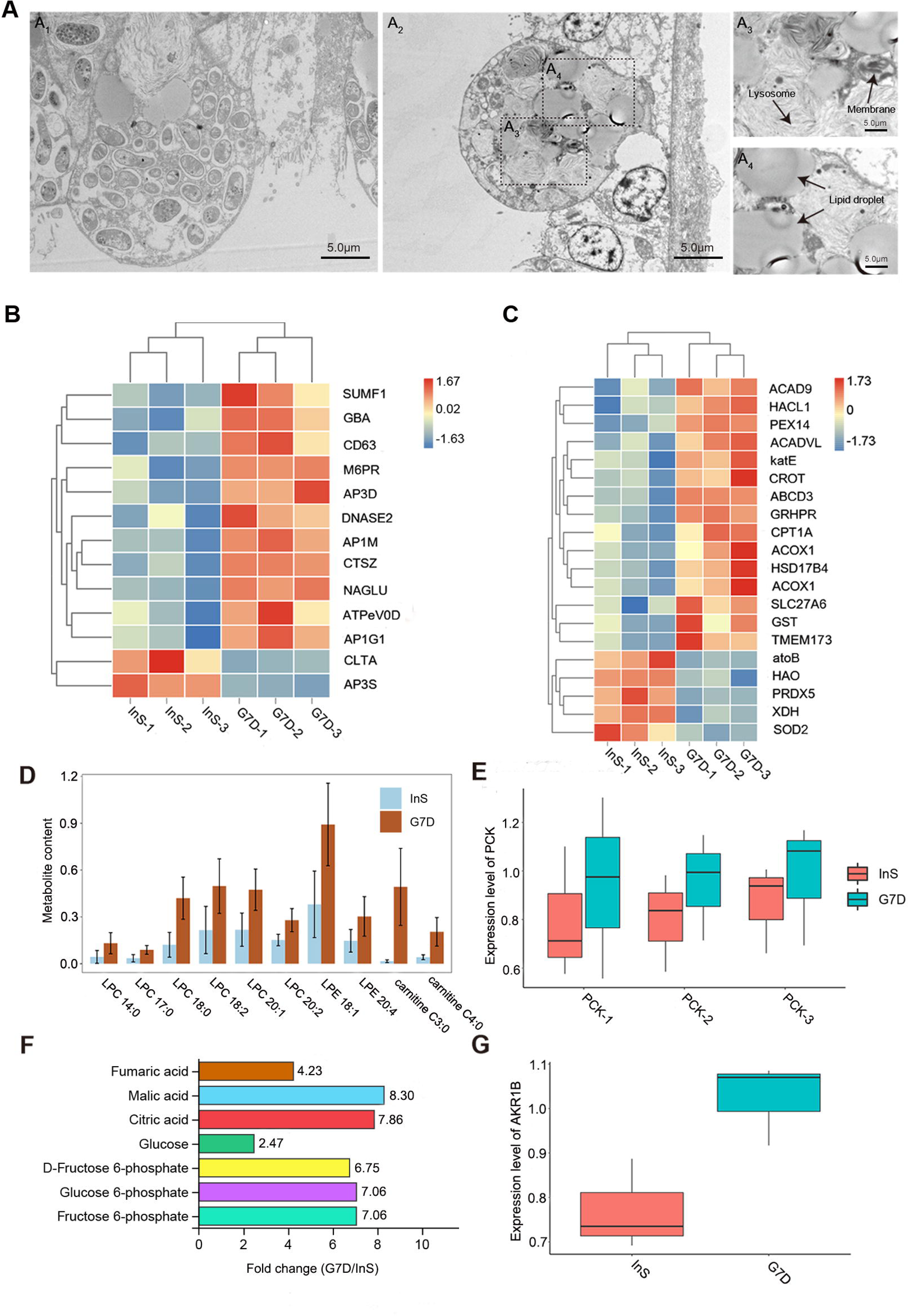
Metabolic remodeling of gill tissue at the early stage of methane deprivation. **(A)** Transmission electron microscope (TEM) images of bacteriocytes in the InS (A_1_) and G7D (A_2_) groups. Methanotrophic endosymbionts were found to be greatly reduced under 7-day methane deprivation, while some digested debris was observed in the augmented lysosomes (A_3_). Large volumes of lipid droplets were also observed in the bacteriocytes shortly after 7-day methane deprivation (A_4_). **(B)** Heatmap of differentially expressed proteins involved in lysosome-mediated digestion in the G7D group in comparison with the InS group. The majority of the differentially expressed proteins were up-regulated shortly after methane deprivation, showing augmentation of lysosome-mediated digestion. Up-regulated proteins are labeled in red, while down-regulated proteins are labeled in blue. **(C)** Heatmap of differentially expressed proteins involved in fatty acid metabolism in the G7D group in comparison with the InS group. The majority of the differentially expressed proteins were up-regulated shortly after methane deprivation, showing augmentation of the β-oxidation of fatty acids. **(D)** Increases in the relative abundance of lysophosphatidylcholine (LPC), lysophosphatidylethanolamine (LPE), and carnitine in G7D group in comparison with InS group as detected by liquid chromatography–mass spectrometry (LC-MS) (data given as mean ± SD). The results confirmed the accumulation of some intermediates of fatty acid oxidation after methane deprivation. **(E)** Expression patterns of three phosphoenolpyruvate carboxykinases (PCKs) homologous in the G7D and InS groups as revealed by the proteome (data given as the mean ± SD). All three enzymes increased shortly after methane deprivation, implying the promotion of gluconeogenesis. **(F)** Fold changes of the relative abundances of carbohydrates in the G7D group compared to the InS group as detected by gas chromatography–mass spectrometry (GC-MS). All gluconeogenesis-related carbohydrates increased markedly in mussels deprived of methane. **(G)** Expressional increase of aldehyde reductase (AKR1B) protein in the G7D group in comparison with the InS group. The increase of AKR1B protein implied the promotion of glycerol synthesis.

It was also noteworthy that the continuous digestion of symbionts might led to hypernutrition in the gill cells, as evidenced by the promotion of host pyruvate metabolism and the increase in gluconeogenesis- and glycerol synthesis-related proteins. In detail, three phosphoenolpyruvate carboxykinase (PCK) proteins, rate-limiting enzymes in gluconeogenesis, were dramatically up-regulated in the G7D group (Fig. 2E). Several gluconeogenesis-related carbohydrates, including glucose, glucose 6-phosphate, fructose 6-phosphate, pyruvic acid, malic acid, and citric acid, were also significantly accumulated in the metabolome of the G7D group (Fig. 2F). Similarly, aldehyde reductase (AKR1B) protein, a key enzyme involved in glycerol synthesis, was also found to be up-regulated in the G7D group (Fig. 2G); this was accompanied by more newly synthesized lipid droplets in bacteriocytes (Fig. 2A). These findings also suggested that the mussel host could further store extra nutrition obtained from the lysosome-mediated digestion, which are necessary in case of prolonged methane deprivation and symbiont loss.

### 3.3. Apoptosis of bacteriocytes highlighted the developmental changes of the gill during the middle stage of methane deprivation

As methane deprivation and symbiont depletion continued, the morphology and cell types of gill filaments began to change noticeably. In particular, the majority of the bacteriocytes disappeared, with only a small number retained at the third month of laboratory methane deprivation. The gill filaments were occupied by ciliated cells or gill cells with microvilli (Fig. 3A), which is commonly observed in non-symbiotic shallow-water mussels and suggested that the mussel host adjusted the development of gill tissue in response to the continued methane deprivation and symbiont depletion. We then examined the proteomics data of the G3M group to screen possible genes and processes guiding these developmental changes. Because the long-term maintenance of symbiotic associations under laboratory conditions remains a common challenge for research in Bathymodiolinae mussels (Hiebenthal et al., 2021), mussels from the InS group and G7D group served as control groups here to exclude the potential influence of the laboratory cultivation system and characterize the key molecules in detail. Consequently, two caspase proteins (caspase 2 and caspase 7) were found to be significantly up-regulated in the G3M group in comparison with mussels during the early stage of methane deprivation (G7D group). Among them, caspase 7 was even found to be markedly down-regulated in the G7D group in comparison with the symbiotic mussels (InS group). These results implied the participation of cell apoptosis in the abolition of bacteriocytes from the early stages of symbiont loss (Fig. 3B). Consistent with this, some of the remaining bacteriocytes in the G3M group were found to have undergo apoptosis via TEM analysis, exhibiting a shrunken cellular size, irregular cell nuclei, condensed chromatin, and phagocyte-mediated phagocytosis (Fig. 3C). In addition, KEGG and GO enrichment analysis showed that biological processes involved in cell proliferation and differentiation were significantly up-regulated in the G3M group compared to the InS group (Fig. S3C). Furthermore, the signaling pathways (such as the MAPK signaling pathway and the VEGF signaling pathway) and proteins (9 out of 10) involved in cell proliferation and differentiation, including neurofibromin 1 (NF1), epidermal growth factor receptor (EGFR), dual specificity protein phosphatase (DUSP1 and DUSP3), cytosolic phospholipase A2 (PLA2G4), sphingosine kinase (SPHK), serine/threonine-protein phosphatase 2B catalytic subunit (PPP3C), and prostaglandin-endoperoxide synthase 2 (PTGS2), were significantly up-regulated in the G3M group in comparison with the InS control group (Fig. 3D), which is consistent with the observation that bacteriocytes were replaced by ciliated cells in the G3M group (Fig. 3A).

**Figure 3.**
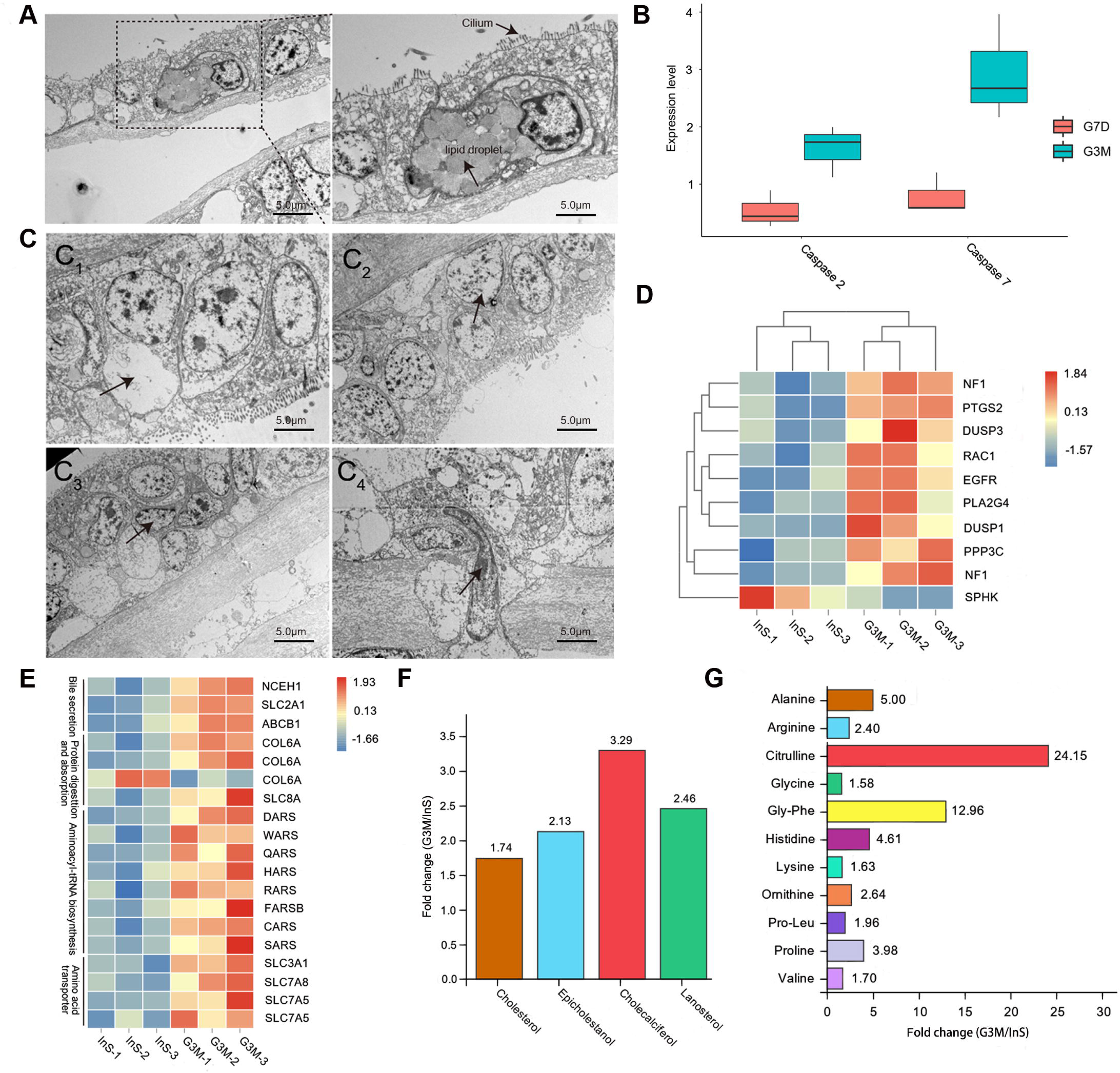
Developmental and metabolic changes of gill tissue at the middle stage of methane deprivation. **(A)** Transmission electron microscope (TEM) images of gill tissue in the G3M group. Few bacteriocytes were retained while most were replaced by ciliated cells in gill tissue at this stage. Scale bar: 5.0 μm. **(B)** Increases in the expression of two caspases (caspase2 and caspase 7) in the G3M group in comparison with the G7D groups as revealed by the proteome (data given as the mean ± SD). The results implied the participation of cell apoptosis in response to methane deprivation and symbiont loss. **(C)** TEM imaging revealed the apoptosis of bacteriocytes with distinct morphological features, including cell shrinkage (C_1_), irregular morphology of the nucleus (C_2_), condensed crescent-shaped chromatin (C_3_), and phagocyte-mediated phagocytosis (C_4_). Scale bar: 5.0 μm. The results confirmed the participation of cell apoptosis in reshaping the morphology of gill tissue. **(D)** Heatmap of differentially expressed proteins involved in cell proliferation and differentiation in the InS group and the G3M group. The majority of the differentially expressed proteins were up-regulated in the G3M group, indicating developmental changes in gill tissue. **(E)** Heatmap of differentially expressed proteins involved in the oxidation of lipids and amino acids in the G3M group compared to the InS group. The results implied augmentation in the turnover and transition of lipids. **(F)** Fold changes of the relative content of steroid metabolites in the G3M group compared to the InS group as identified by the gas chromatography–mass spectrometry (GC-MS) metabolome. The results implied augmentation in the turnover and transition of sterols. **(G)** Fold changes of the relative abundance of amino acids in the G3M group compared to the InS group as revealed by the liquid chromatography–mass spectrometry (LC-MS) metabolome. The results implied augmentation of the turnover and transition of amino acids.

With the depletion of symbionts and apoptosis of bacteriocytes, the mussels could no longer obtain nutrition from symbionts. In consistency with the conclusion, the host down-regulated the lysosome-mediated symbiont-digestion process (Fig. S3C). Moreover, it was found that biological processes involved in protein digestion and absorption were broadly promoted (Fig. S3C). In addition, proteins (18 out of 19) involved in the turnover and transition of essential metabolites, especially sterols and amino acids, were also promoted. For example, neutral cholesterol ester hydrolase 1 (NCEH1) protein, a key enzyme involved in hydrolyzing cholesterol ester into cholesterol, was significantly up-regulated in the G3M group compared to the symbiotic mussels in the InS group (Fig. 3E). Similarly, host genes involved in amino acid transportation and cycling, such as collagen and solute carrier family members (SLC3A1 and SLC7A8) and aminoacyl-tRNA synthetases, were also significantly up-regulated compared to the InS group (Fig. 3E). The up-regulation of these proteins suggested that the host was promoting the turnover process of essential metabolites to satisfy its nutritional needs after prolonged methane deprivation and in the absence of symbionts. In accordance with the proteomics result, increases of cholesterol, lanosterol, epicholestanol, cholecalciferol, and other sterol intermediates were observed in the G3M group, in comparison with the InS control group (Fig. 3F). Similarly, concentrations of some essential amino acids (such as arginine, proline, glycine, valine, ornithine, and alanine), proteolytic intermediates (such as Pro-Leu and Gly-Phe), and by-products of amino acid biodegradation (such as citrulline), were also found to be increased in the G3M group compared to the InS group (Fig. 3G).

### 3.4. Enhanced filter-feeding function facilitated long-term survival of mussels without methane supply and methanotrophic symbionts

As mentioned above, the methanotrophic deep-sea mussels could survive for years (more than 300 days in the present study) after methane deprivation-induced symbiont loss. This finding raised the question of how the mussel hosts adjusted the metabolism and function of gill tissue after the complete loss of bacteriocytes and the replenishment of ciliated cells. It is speculated that the mussels could continue down-regulate their energy-consuming processes to conserve nutrition, which is supported by the continuous down-regulation of immune-related processes such as the bacterial invasion of epithelial cells and endocytosis (Fig. S3ABD). Furthermore, glucose metabolism (including glycolysis, the TCA cycle, and oxidation phosphorylation) and amino acid metabolism increased significantly in the gill tissue of the G10M group in comparison with the InS control group (Fig. S3D). For example, key enzymes in glucose metabolism, such as aldose 1-epimerase (galM), glucose-6-phosphate 1-epimerase, 6-phosphofructokinase 1 (pfkA), fructose-bisphosphate aldolase (ALDO), pyruvate carboxylase (PC), citrate synthase (CS), aconitate hydratase (ACO), 2-oxoglutarate dehydrogenase (OGDH), succinyl-CoA synthetase beta subunit LSC1/2, succinate dehydrogenase, fumarate hydratase (fumA), malate dehydrogenase (MDH1), NADH dehydrogenase, ubiquinol-cytochrome c reductase CYC/QCR7, and cytochrome c oxidase, were found to be markedly promoted in mussels that had been decolonized for 10 months (Fig. 4A, B). While the deep-sea mussels promoted glucose metabolism, we also noted that the NCEH1 protein, a key enzyme in fatty acid oxidation, was continuously up-regulated in the G10M group (Fig. S5D). In addition, proteins involved in the oxidation of amino acids were also up-regulated, which was confirmed by the sharp decrease in glutamine, glutamate, alanine, and ornithine in the metabolomics analysis; this could serve as alternative means to obtain energy under conditions of long-term starvation (Fig. 4B). Consistent with the promotion of the oxidation of acid and amino acids, four enzymes involved in mitophagy and protection from oxidative damage, including TBC1 domain family member 15 (TBC1D15), serine/threonine-protein phosphatase (PGAM5), optineurin (OPTN) and Ras-related protein M-Ras (MRAS), were also found to be promoted in the G10M group compared to the symbiotic mussels in the InS group (Fig. 4C). Accompanied by the promotion of energy production, it was noted that proteins involved in cytoskeleton rearrangement and cilium movement began to be up-regulated in the G10M group (in comparison with the G3M group, Fig. 4D). The recovery of cilium movement was consistent with the observation that the host replaced bacteriocytes with ciliated cells (Fig. 3A), which plays important roles in the filter-feeding of the gill in obtaining nutrients from the surrounding environment.

**Figure 4.**
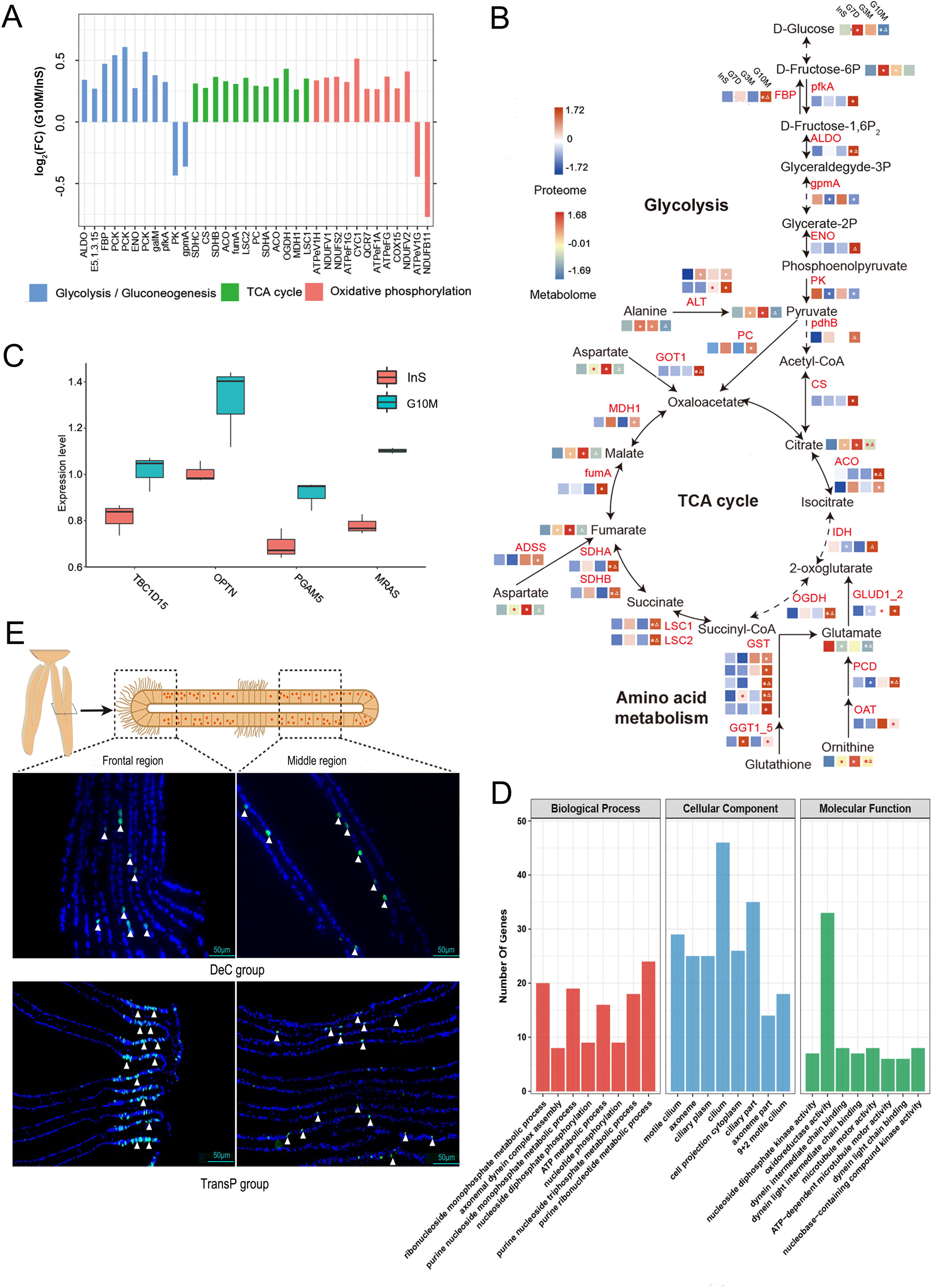
Phenotypic changes of gill tissue under long-term methane deprivation. **(A)** Fold changes of the differentially expressed enzymes involved in glycolysis, the TCA cycle, and oxidation phosphorylation in the G10M group compared to the InS group. The majority of the differentially expressed proteins were up-regulated in the G10M group, indicating the promotion of glycolysis, the TCA cycle, and oxidative phosphorylation. **(B)** Overview of glycolysis, the TCA cycle, and amino acid metabolism under the methane deprivation assay. Changes in the relative abundances of metabolites and proteins are illustrated in the heatmap. Columns in the heatmap (from left to right) represent the relative abundances in the InS, G7D, G3M, and G10M groups. Asterisks represent the significant differences in the G10M group in comparison with the InS group. Triangles represent the significant differences in the G10M group in comparison with the G3M group. **(C)** Box plot showing the four up-regulated enzymes involved in mitophagy in the G3M group compared to the InS group. The results implied the participation of mitophagy in response to methane deprivation and symbiont loss. **(D)** Gene Ontology (GO) enrichment analysis of differentially expressed proteins in the G10M group compared to the G3M group. The top eight most significant GO terms involved in the biological process, cellular component, and molecular function are shown. The processes involved in cytoskeleton rearrangement and cilium movement were enriched in the differentially expressed proteins of the G10M group. **(E)** 5-ethynyl 2′-deoxyuridine (EdU)-labeling assay of gill tissue under long-term methane deprivation. Mussels in the G10M group were subjected to EdU treatment to label newly generated cells under either *ex-situ* (in laboratory recirculating system) or *in-situ* conditions (transplanted back to the cold seep). Newly synthesized DNA was incorporated with EdU and visualized in green fluorescence. Cell nuclei were stained by DAPI (in blue). The gills of the G10M group retained cell proliferation ability, and more intensive signals were observed in mussels transplanted back to cold seeps.

### 3.5. Robust cell proliferation highlighted potential resilience of mussels under methane reduction and deprivation

While the mussels managed to survive through the long-term methane deprivation at the expense of losing all symbionts and changing phenotype of gill tissue, we then asked whether these decolonized mussels could recover from these changes once the methane are accessible again. Since the cell proliferation of gill tissue could guarantee a continuous supply of new cells to mount a rapid and dynamic response to emerging environmental changes (e.g., either prolonged methane deprivation or the rejuvenation of methane), we then conducted the EdU incubation assay with the decolonized mussels to label cell proliferation and to indicate the potential resilience of mussels under methane deprivation. It turned out that signs of proliferating cells could be observed in the G10M mussels, suggesting the proper functioning of gill tissue (Fig. 4E, the DeC group). Moreover, a more intensive signal of cell proliferation was observed in the mussels that were returned and transplanted back to the seepage region or the low-methane region of the cold seep (the TransP group). Although no signs of recolonized symbionts were observed, the enhanced cell proliferation indicated that the host generated more cells for the rejuvenation of methane (Fig. 4E).

### 3.6. Sterol metabolism as a direct cue in triggering mussel’s responses to methane deprivation

As demonstrated above, the deep-sea mussel has altered its global metabolism, cell function and tissue development dynamically during methane deprivation. However, the direct cues that trigger the mussel’s responses to methane deprivation remained unknown. Considering that the proteins responsible for the phenotypic changes in gills could share a coordinated expression pattern across samples and thus constitute co-expression networks, we then conducted the weighted gene co-expression network analysis (WGCNA) to identify the key transcription factors or signal transducers in the protein network guiding the phenotypic changes of gill. Consequently, a total of 18 co-expression modules were obtained for the proteome of deep-sea mussels, among which mod2, mod3, mod5, mod13, and mod17 were of particular interest as they contained the majority of differentially expressed proteins. For example, mod2, mod5, and mod13 contained differentially expressed proteins in the G7D/G4W, G3M, and G10M groups in comparison with the InS group; mod17 contained differentially expressed proteins in the G3M group in comparison with the G7D group; and mod3 contained differentially expressed proteins in the G10M group in comparison with the G3M group (Fig. S7). Noticeably, 42 differentially expressed proteins from the G7D/G4W group (in comparison with the InS group) were found to be highly interconnected (as a co-expression network) within mod2, and were involved in cell metabolism, signal transduction, and transcription regulation. Among these proteins, six proteins, including nuclear receptor-binding factor 2 (NRBF2) and ATP-dependent RNA helicase A (DHX9), were annotated as transcription regulators (Fig. 5A). In particular, NRBF2 protein, a crucial regulator in starvation-induced autophagy, showed the highest interconnectivity degree with the remaining differentially expressed proteins (connecting 33 of these proteins), which might serve as the hub regulators during the early stage of methane deprivation. In addition to the NRBF2 protein, HSD17B4, a crucial enzyme in sterol and steroid hormone metabolism, was also found to have a high interconnectivity degree with the differentially expressed proteins of mod2 (Fig. 5A), indicating its role in regulating methane deprivation-induced phenotypic changes. In accordance with these findings, two additional sterol-related signaling pathway proteins (24-hydroxycholesterol 7 alpha-hydroxylase and nuclear receptor coactivator 6) were also found to be differentially expressed in the G7D group, accompanied by the drastic alterations of sterol-related metabolites in gill tissue. These results collectively suggest that the sterol-related signaling pathway may play a crucial role in initiating the metabolic, function, and development changes of gill tissue shortly after methane deprivation (Fig. 5B).

**Figure 5.**
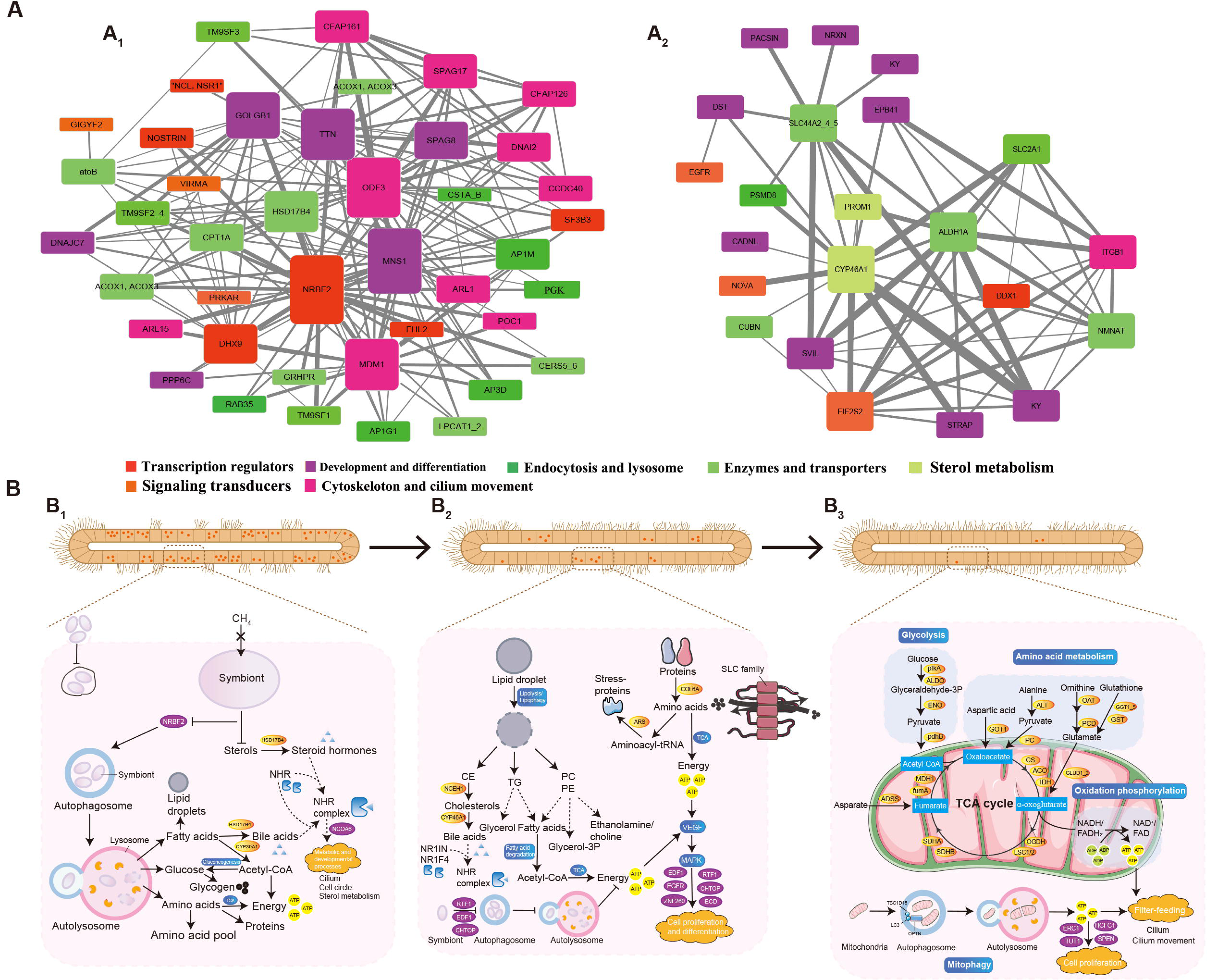
Regulatory networks of deep-sea mussels under methane deprivation. **(A)** Regulatory networks of gill tissue under methane deprivation constructed using weighted gene co-expression network analysis (WGCNA). About 42 differentially expressed proteins (nodes in the network) in the G7D/G4W groups in comparison with the InS group were found to be interconnected in mod2 (A_1_), constructing the hub regulatory networks at the early stage of methane deprivation. NRBF2 and HSD17B4 had a high degree of interconnectivity with the differentially expressed proteins in mod2, suggesting the participation of sterol and steroid hormone metabolism in regulating the phenotypic changes of gill tissue at early stages of methane deprivation. Similarly, a total of 22 differentially expressed proteins in the G3M group in comparison with the InS group were interconnected in mod5 (A_2_), constructing the hub regulatory networks at the middle stage of methane deprivation. Among the hub genes, CYP46A1 protein was also up-regulated in the G3M group and exhibited the highest interconnectivity with other proteins, suggesting the ongoing contribution of the sterol-related signaling pathway after losing all symbionts under methane deprivation. Proteins with similar function or in same biological process are labeled using the same color. Interconnectivity degree of corresponding protein is represented by the node height. Calculated weight between two corresponding proteins is represented by the width of edge. **(B)** Schematic map showing the metabolic, developmental and functional changes of gill tissue at early-stage (B_1_), middle-stage (B_2_), and long-term (B_3_) methane deprivation. Purple boxes indicate transcription regulators. Yellow boxes indicate enzymes. Blue boxes indicate metabolic processes. Solid lines or arrows show the reactions catalyzed by differentially expressed proteins identified by the proteome. Dotted lines or arrows show the reactions catalyzed by non-significantly differentially expressed proteins identified by the proteome.

Similarly, a total of 14 transcription regulators, including endothelial differentiation-related factor 1 (EDF1), chromatin target of PRMT1 protein (CHTOP), RNA polymerase-associated protein RTF1 homolog (RTF1), protein ecdysoneless homolog (ECD), zinc finger protein 260 (ZNF260), and EGFR, were identified in mod5 and mod17, modulating cell metabolism, development, and differentiation processes during middle-stage methane deprivation (Fig. S8A). It was interesting to note that the cholesterol 24-hydroxylase (CYP46A1) protein, another gene involved in sterol turnover and the biosynthesis of steroid hormone, was also up-regulated in the G3M group and exhibited the highest interconnectivity with other proteins, suggesting the ongoing contribution of the sterol-related signaling pathway to the gill tissue even after losing all symbionts under methane deprivation (Fig. 5A). Comparatively, four transcription regulators, including ELKS/Rab6-interacting/CAST family member 1 (ERC1), host cell factor 1 (HCFC1), protein split ends (SPEN), and translation regulators speckle targeted PIP5K1A-regulated poly(A) polymerase (TUT1), were found to be differentially expressed in the G10M group, potentially influencing the expression of 59 proteins that were involved in cell metabolism and cilium movement, therefore constituting the core modulatory network under long-term methane deprivation (Fig. 5B and Fig. S8B, C).

## 4. Discussion

Being a new unconventional alternative energy, the exploration and exploitation of methane hydrates has become one of the hottest topics in the field of energy, engineering, and environment. While there are still many technical challenges to safely and economically mine the methane hydrates that are buried above or under the seafloor, environmental impact assessment of the methane hydrate mining is another major concern to be considered (Boswell, 2009). Here, we assessed the long-term influences of simulated methane hydrate mining to the methanotrophic deep-sea mussel *G.platifrons*, a dominate and endemic mollusc in cold seeps of west Pacific, with laboratory methane deprivation assay and *in situ* transplantation assay. Our results collectively show the drastic changes in the metabolism, function and development of mussel gill during the 10-month methane deprivation, and provide a holistic data to understand the endurance and potential resilience of deep-sea mussel to methane hydrate reduction and deprivation, which caused by mining.

Unlike their shallow-water relatives, deep-sea mussels have less access to photosynthetic carbon, and thus rely mostly on chemosynthetic carbon to survive. While deep-sea mussels can harvest nutrition from their chemosynthetic endosymbionts via either “milking” or “farming” (Kádár et al., 2008; Streams et al., 1997), their nutritional balance breaks down as soon as the number of endosymbionts or the amount of environmental chemosynthetic resources (e.g. methane for methanotrophic symbionts) decrease. As expected, the metabolic processes of methanotrophic hosts were found to be altered soon after methane deprivation. In particular, the deep-sea mussels digested their endosymbionts rapidly and in massive quantities to meet their nutritional needs via a lysosome-mediated process, which agrees with previous studies on deep-sea mussels (Détrée et al., 2019; Sun et al., 2022; Wang et al., 2019), and which is often observed in other symbiotic associations (Bright et al., 2000; Streams et al., 1997; Volland et al., 2018). It is also noticeable that immune processes that could benefit the horizontal acquirement of symbionts, including bacterial invasion of epithelial cells and endocytosis (Tame et al.), were repressed by the gill since the methane deprivation. In addition, promotion in the lysosome-mediated digestion of symbionts also ceased when the methane deprivation continued and fewer symbionts were left. Although the down-regulation of immune processes might influence the recolonization of symbionts, it is yet energy-saving in the long-term starvation caused by methane deprivation.

What is more interesting is that the deep-sea mussels can survive for years even after their complete loss of symbionts as observed in our and other studies (Tame et al.), highlighting the drastic endurance of deep-sea mussels to the inhabitable environment. To sustain the aposymbiotic life, the mussels were found to gain energy by promoting the turnover process of essential metabolites, as evidenced by the upregulation of protein digestion and absorption, and turnover of sterol and amino acids in G3M and G10M group. Additionally, programmed cell death of bacteriocytes is found massively activated, serving as an extra source of nutrients. Although the digestion of self components is often observed across species during starvation (Chera et al., 2009; Kawabata and Yoshimori, 2016), we also found that the mussels might reallocate energy to sustain the proliferation of ciliated cells and enhance the filter-feeding function of the gill after their complete loss of symbionts and despite of having a tight energy budget. In addition, intensive signals of DNA replication along the gill filaments were observed in our EdU labeling assay and confirmed the widespread of stem-like cells in gills. The stem-like cells interspersed along the gill were believed to be intercalary cells and could serve as important source of the new born ciliated cells (Piquet et al., 2022). The gains and losses of different gill cells could also be observed in the thiotrophic deep-sea mussel *Bathymodiolus azoricus* after a 2-months starvation (Piquet et al., 2022), showing the robust phenotypic plasticity of gill in adult mussels. These phenotypic changes provide an alternative method for the deep-sea mussel to obtain nutrients by taking them directly from the external environment, which is the ancestral method that non-symbiotic shallow-water mussels use (Page et al., 1990).

Since the intensive signals of cell proliferation and renewal of ciliated cells confirmed the drastic endurance of deep-sea mussels to methane deprivation, we then questioned on whether these long starved and decolonized mussels could recover from the methane reduction or deprivation when methane are accessible again. To answer this question, we have transplanted the decolonized mussels back to the deep-sea and retrieved five days later (TransP group). It surprisingly turned out that more signals of proliferating cells could be observed in the transplanted mussel gills, implying rejuvenation of gill function and development. For the thiotrophic Bathymodiolonae mussels, it has shown that the symbiotic associations could recover gradually in some once sulfide becomes available once more (Hiebenthal et al., 2021; Kádár et al., 2005). However, the bacterial signals yet were still not observed in this study, suggesting that the recolonization of symbionts may need to take more time. It should also be noticed that the sulphide deprivation in above study is only about 30 days while the recovery took about 15 days (Kádár et al., 2005). Therefore, the endurance thresholds of deep-sea mussel in methane deprivation to permit symbiont recolonization should be further determined. On the other hand, it is also would be interesting to determine the long-term impact (especially the transgenerational effects) of symbiont depletion on the deep-sea mussels, as there have been some reports on the loss or reduction of symbiosis in the bathymodiolin mussel and other deep-sea bivalves (Rodrigues et al., 2015; Sibuet and Olu, 1998). Nevertheless, the endurance and resilience of deep-sea mussels to methane reduction and deprivation is believed to be a major issue from both the biological and ecological aspect, which could facilitate the adaptation and widespread of deep-sea mussels in the fluctuating environment of deep-sea and provide baseline information to protect them in future methane hydrate mining.

When evaluate the ecological impact of deep-sea mining, including methane hydrate mining, it is important to ascertain the biomarkers of megafauna to these environmental stresses induced by the mining. To date, only a few studies have been carried out to identify the biomarkers of megafauna to deep-sea mining and majority of them were conducted using the shallow-water proxies (Carreiro-Silva et al., 2022; Galgani et al., 2005; Marassi et al., 2023; Pinheiro et al., 2019; Pinheiro et al., 2021).

Although the biological processes of shallow-water proxies may resemble these of the deep-sea megafauna, the stress responses yet could be species-specific, demonstrating the necessity of studies with local species (Mestre et al., 2019). With insightful data from the local megafauna such as deep-sea mussels, the pelagic helmet jellyfish, deep-sea crustaceans and some deep-sea corals, it is clearly showed that some immune-related processes, the antioxidant system, tissue regeneration- or wound repair-related genes and cellular metabolism could serve as biomarkers of heavy metal, sediment plume and oil stresses (Auguste et al., 2016; Bebianno et al., 2018; Company et al., 2019; Deleo et al., 2018; Martins et al., 2017; Stenvers et al., 2023; Zhou et al., 2021). Here, our multi-omics data clearly showed that the methane deprivation could be a vital stress to the mussel holobionts considering the drastic alternations in the metabolism, function, and development of gill and the rapid depletion of symbionts. Meanwhile, the stress responses of deep-sea mussel to methane deprivation could be divided into three distinct stages in accordance with the remaining abundance of symbionts (Fig. 5B), and therefore could indicate the health state of deep-sea mussels during methane hydrate mining.

While the biomarkers reflect the global responses of deep-sea mussels to methane deprivation, we further asked if there were some leading metabolites, genes or pathways that control the dynamic responses during long-term methane deprivation. Here, we found that the sterol-related signaling pathway and nuclear receptors might play important roles in regulating the biological responses of gill tissue to the methane deprivation. Noticeably, the sterol metabolism of mussel hosts is reported to rely greatly on that of their symbionts (Takishita et al., 2017), and can be altered in compensatory ways after symbiont depletion (Sun et al., 2022). It is therefore suggested that the drastic changes in the host biological processes might be triggered by the shortage of symbiont-deprived sterol intermediates, a cascade reaction induced by the methane deprivation. The cascade reaction also highlights the participation of symbiont in the phenotypic plasticity of symbiotic organs, which can also be evidenced by other studies and data from our single-cell transcriptome analysis. For example, it is noted that the epithelial cells of deep-sea mussels can become hypertrophic and lose microvilli and cilia once colonized by symbionts (Franke et al., 2021). In addition, the newly formed bacteriocytes of adult mussels also lose microvilli after being colonized by the symbionts of adjacent bacteriocytes (Wentrup et al., 2014). In our single-cell transcriptome study, we showed the participation of the sterol-related signaling pathway in the differentiation and maturation processes of mussel bacteriocytes (Chen et al., 2023). It is also noteworthy that the symbionts may control the function and development of host symbiotic organs in other symbiotic associations (Coon et al., 2017; Heath-Heckman et al., 2016; Nyholm and Mcfall-Ngai, 2021). Although the detailed mechanisms beneath the symbiont-mediated phenotypic plasticity of symbiotic organs still need to be further studied in future, these findings collectively reconfirmed that the mussel hosts could reshape its metabolism, function and development of gill based on direct cues from symbionts.

There were some limitations to the current study. For example, while the global metabolism and gene expression of endosymbionts could be influenced by depressurization stress (Chen et al., 2021), it is still challenging to cultivate deep-sea mussels in pressurized laboratory systems and to conduct simulations in deep-sea, which complicated the interpretation of the results. In addition, the maintenance of symbiotic associations under laboratory conditions is also challenging for deep-sea holobionts (Hiebenthal et al., 2021; Sun et al., 2022), which therefore complicates the choice of a proper control group during long-term laboratory cultivation. Alternatively, deep-sea mussels collected from the seepage region in an isobaric and isothermal way were employed as the control group to represent the natural state of symbiotic association. It is noteworthy that these limitations also hamper laboratory investigations on the resilience of deep-sea mussels to methane reduction and deprivation, which therefore should be conducted in deep-sea under *in situ* conditions in future.

## 4. Conclusions

Taken together, the present study depicted the phenotypic changes of gill tissue in response to simulated methane hydrate mining based on the integrated multi-omics data. The observed dynamic remodeling of the cell metabolism, development, and function of gill tissue support the long-term survival of deep-sea mussels even after the loss of all endosymbionts, which advantages the dispersal and persistence of Bathyodiolinae mussels in the harsh environment of the deep sea and provides essential information for the deep-sea methane hydrate exploration and exploitation.

## Acknowledgments

We thank the crew members of the R/V Kexue for their assistance in sample collection and the laboratory staff for continuous technical advice and helpful discussions. We also thank Professor Guowang Xu and Professor Xinjie Zhao for their assistance in the metabolome analysis. We also thank Prof. Qiang Lin from the South China Sea Institute of Oceanology, Chinese Academy of Sciences for insightful comments and suggestions that improved the manuscript.

## Funding

The present study was supported by the National Natural Science Foundation of China (42030407, 42106134, 42076091, and 42221005), the Key Research Program of Frontier Sciences (ZDBS-LY-DQC032), Laoshan Laboratory (No. 2022QNLM030004), the Strategic Priority Research Program of the Chinese Academy of Sciences (XDA22050303 and XDB42020401), and the Key Deployment Project of the Center for Ocean Mega-Science, CAS (COMS2020Q02).

## Author contributions

M.L. and H.C. participated in the study design and data interpretation. H.C., M.W., S. Z., and C. Lian helped in the sample collection. M.L. conducted the majority of the experimental work and data analysis and drafted the manuscript. Y.L., M.W., S. Z., L.Z., H.Z., and H.W. assisted with the interpretation of the results. L.C. surveyed environmental parameters during the cruise. H.C. and C. Li conceived the study, coordinated the experiments, and helped draft the manuscript. All authors gave their final approval for publication.

## Competing interests

The authors declare no competing interests.

## Data accessibility

All data needed to evaluate the conclusions in the paper are presented in the paper and/or the supplementary materials. All proteomic raw data have been deposited into the iProX Consortium (https://iprox.org/) with the project ID IPX0004524000. All metabolome raw data have been deposited into the MetaboLights database (https://www.ebi.ac.uk/metabolights/) with the identifier MTBLS4998. All meta-transcriptome sequencing data have been deposited in the NCBI BioProject database (https://www.ncbi.nlm.nih.gov/bioproject/) under accession no. PRJNA912760.

## Supplementary Figures

**Fig. S1. Gnome mapping rate of endosymbionts in mussels of the InS and InS_At (*in situ* antibiotics treatment) groups.**

**Fig. S2. Overview of gill proteome under methane deprivation.**

**(A)** The relative abundance of the *pmoA* gene in the InS, InS_At (*in situ* antibiotic treatment), G7D, G3M and G10M groups. The DNA content of *pmoA* gene was no longer detectable by qRT-PCR in the G10M group.

**(B)** Relative abundance of the *pmoA* gene in gill tissues of mussels acclimated under 2 MPa pressure (2MPa group) or atmospheric pressure (NP group). The relative abundances of the pmoA gene remained comparable on the first (D1), third (D3) and sixth (D6) days after acclimation.

**(C)** UpSet diagram of the differentially expressed proteins in the G7D, G3M, and G10M groups compared to the InS group. The horizontal bars show the total number of differentially abundant proteins in each group compared to the InS group. The vertical bars show the shared number of differentially abundant proteins between the given groups. Connected dots refer to the groups involved in each intersection.

**(D)** Principal component analysis (PCA) of proteome data of all groups. The proteomic data from decolonized mussels (expect for the G3M group) were clustered distinctly apart from those of symbiotic mussels (InS group).

**(E)** Pearson correlation analysis of the proteome in all groups (red indicates a high correlation, white indicates a low correlation). The results showed well consistency among samples of the same group, verifying the repeatability and reliability of the proteomic data.

**(F)** Venn diagram of differentially expressed proteins in the G7D and G4W groups compared to the InS group. The majority of the differentially expressed proteins were shared between the G7D and G4W groups.

**Fig. S3. Kyoto Encyclopedia of Genes and Genomes (KEGG) enrichment analysis of differentially expressed proteins.**

The significantly enriched KEGG pathways of up-regulated and down-regulated differentially expressed proteins in the G7D group (A), G4W group (B), G3M group (C), and G10M group (D) compared to the InS group are illustrated. The results showed that cell metabolism, cytoskeleton/organelle rearrangement and cell proliferation/differentiation processes were enriched in decolonized mussels.

**Fig. S4. Gene Ontology (GO) enrichment analysis of differentially expressed proteins in the G7D, G4W, G3M, and G10M groups compared to the InS group.**

The top 20 most significant GO terms of the differentially expressed proteins of the G7D group (A), G4W group (B), G3M group (C), and G10M group (D) compared to the InS group are illustrated. The circles from the outside to the inside indicate the classification of GO terms (yellow indicates biological process, blue indicates molecular function, and green indicates cellular component) and the number of background proteins in each classification (longer indicates more proteins). Colors are used to indicate the *p*-values (red indicates greater significance), the numbers of up-regulated (intense purple) and down-regulated (light purple) proteins, and the rich factor value of each classification (each compartment represents 0.1). The results showed that GO terms involved in cell metabolism, cytoskeleton/organelle rearrangement and cell proliferation/differentiation were enriched in decolonized mussels.

**Fig. S5. Overview of metabolome analysis in the InS, G7D, G3M, and G10M groups.**

**(A)** Principal component analysis (PCA) of metabolites identified in the InS, G7D, G3M, and G10M groups by liquid chromatography–mass spectrometry (LC-MS).

**(B)** PCA analysis of metabolites identified in the InS, G7D, G3M, and G10M groups by gas chromatography–mass spectrometry (GC-MS).

**(C)** UpSet diagram of the differentially expressed metabolites in the G7D, G3M, and G10M groups compared to the InS group. The horizontal bars show the total number of differentially abundant metabolites in each methane deprivation treatment group compared to the InS group. The vertical bars show the shared number of differentially abundant metabolites between the given groups. Connected dots refer to the groups involved in each intersection.

**(D)** Expression pattern of neutral cholesterol ester hydrolase 1 (NCEH1) during methane deprivation (data given as the mean ± SD).

**Fig. S6. KEGG enrichment analysis of differentially expressed genes in *in situ* antibiotic treated mussels compared to InS mussels revealed by transcriptome.**

**Fig. S7. Sample expression pattern and Venn diagrams of differentially expressed proteins identified in the G7D, G4W, G3M, and G10M groups and by weighted gene co-expression network analysis (WGCNA).**

**(A)** WGCNA Sample expression pattern revealed a total of 18 co-expression modules (mod1-18).

**(B)** Venn diagram of differentially expressed proteins in the G7D group and in mod2.

**(C)** Venn diagram of differentially expressed proteins in the G4W group and in mod2.

**(D)** Venn diagram of differentially expressed proteins in the mod5, mod17, and in G3M group.

**(E)** Venn diagram of differentially expressed proteins in the mod13, mod3, and in G10M group.

**Fig. S8. Weighted gene co-expression network analysis (WGCNA) of regulatory networks in the mod17, mod3, and mod13.**

The protein regulatory networks in mod17 (A) and mod3 (B) in the G3M group and mod13 (C) in the G10M group as constructed by WGCNA are illustrated.

## Supplementary Tables

**Supplementary Table S1. Differentially expressed genes of *G. platifrons* host and symbionts between the InS_At group and the InS group in the meta-transcriptome analysis.**

**Supplementary Table S2. List of identified proteins in the proteome of *G. platifrons* gills.**

**Supplementary Table S3. Differentially expressed proteins identified in different groups.**

**Supplementary Table S4. Identified metabolites in the metabolome of *G. platifrons* gills.**

**Supplementary Table S5. Differential metabolites identified in different groups.**

## Notes

### Competing Interest Statement

The authors have declared no competing interest.

### Summary of Updates

The introduction and discussion part are extensively revised to clarify the indication role of phenotypic plasiticity of gill in deep-sea mining of gas hydrates.

