## Supplementary figures and images for "Phenotypic Plasticity of Symbiotic Organ Highlight Deep-sea Mussel as Model Species in Monitoring Exploitation of Deep-sea Methane Hydrate"

### Fig. S1

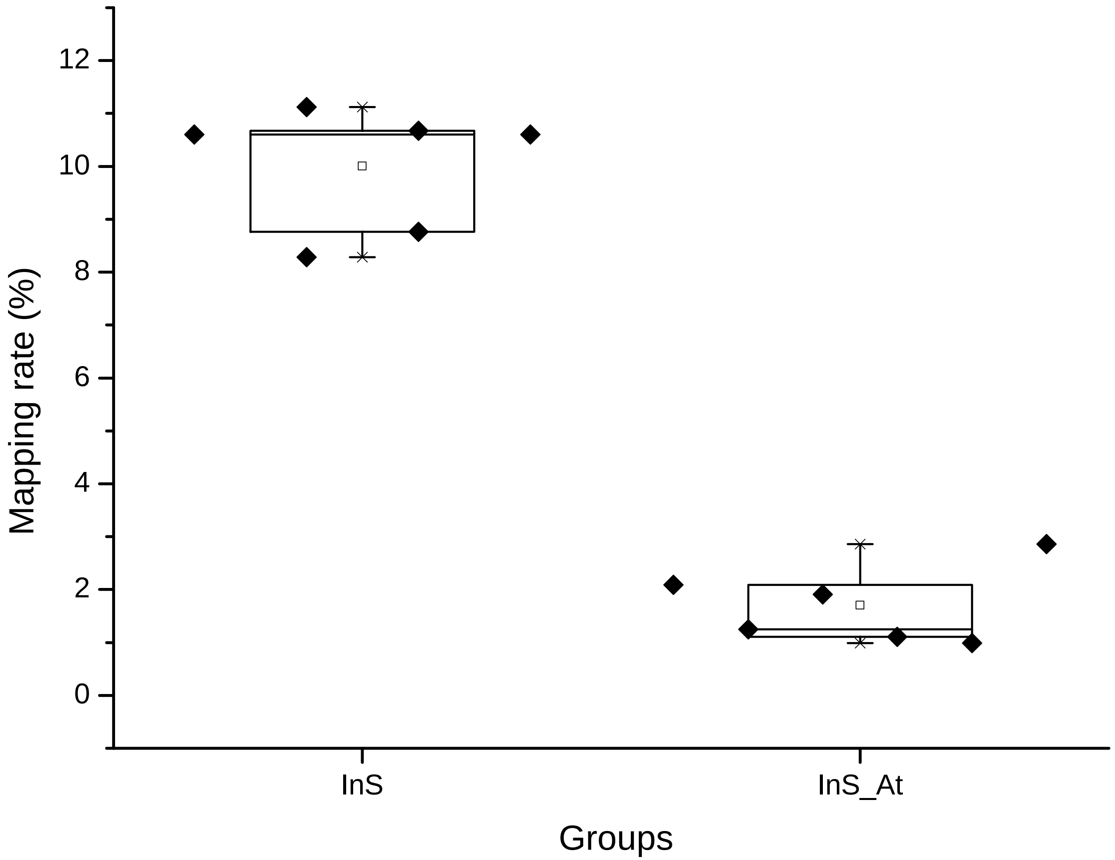

### Fig. S2

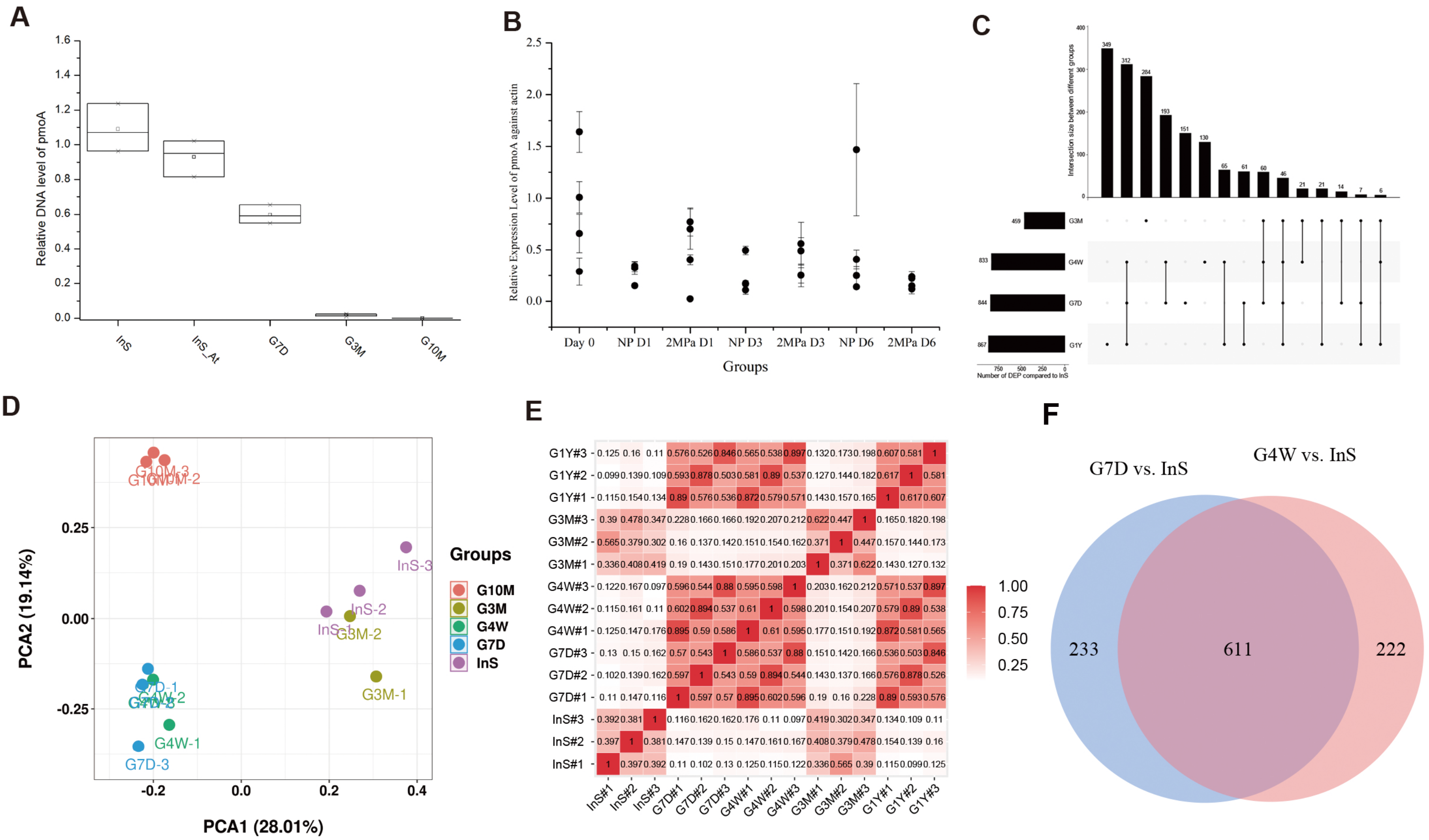

### Fig. S3

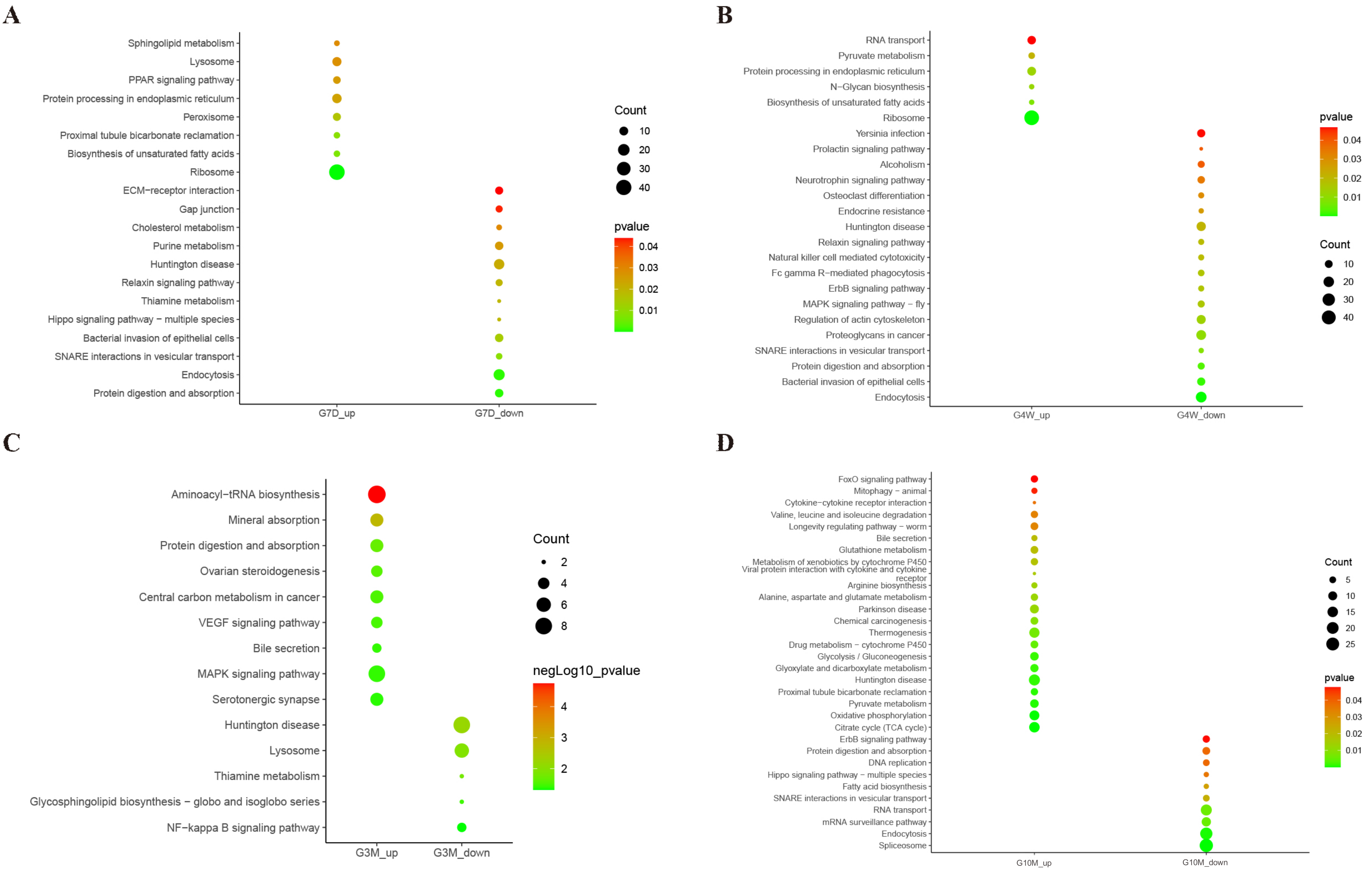

### Fig. S4

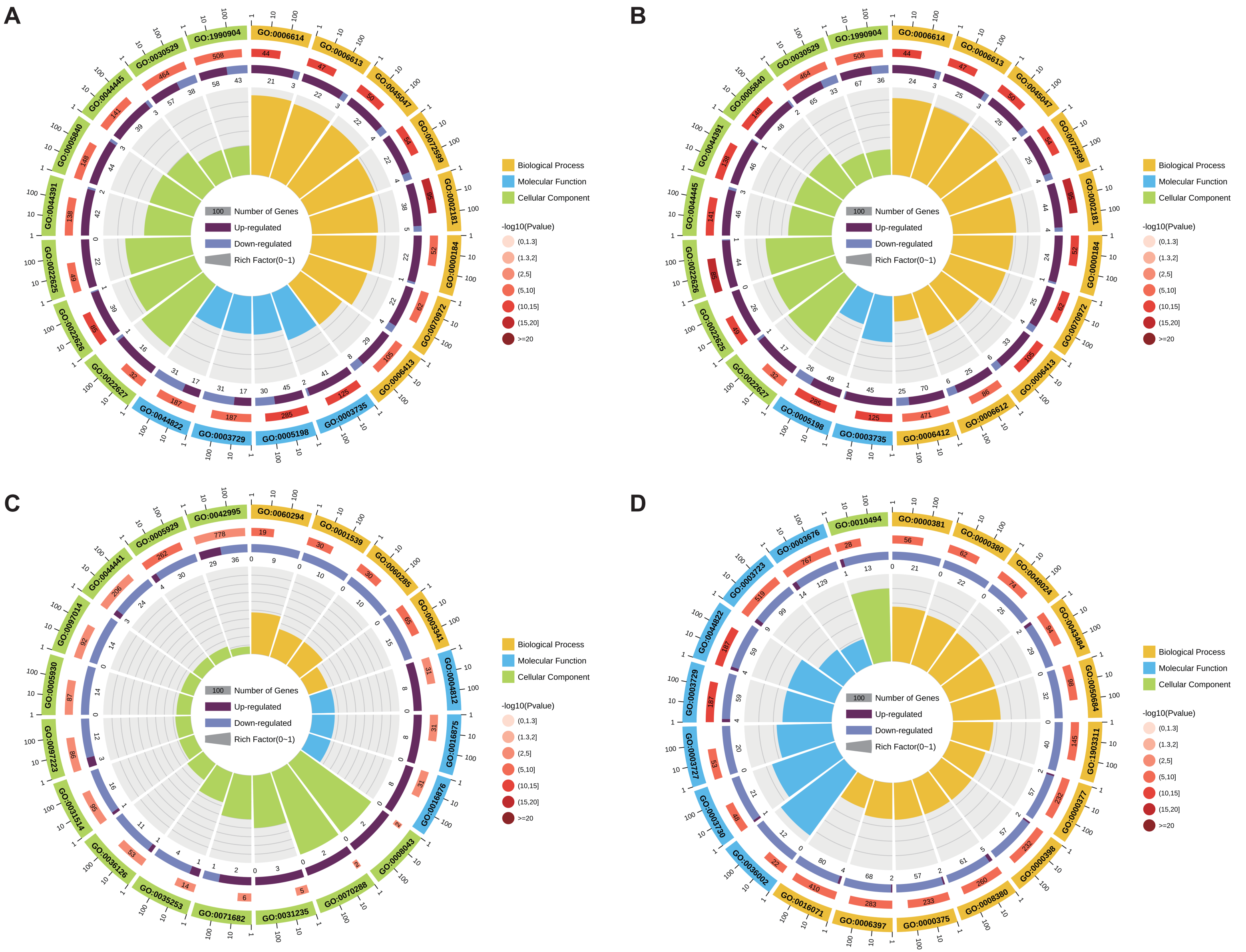

### Fig. S5

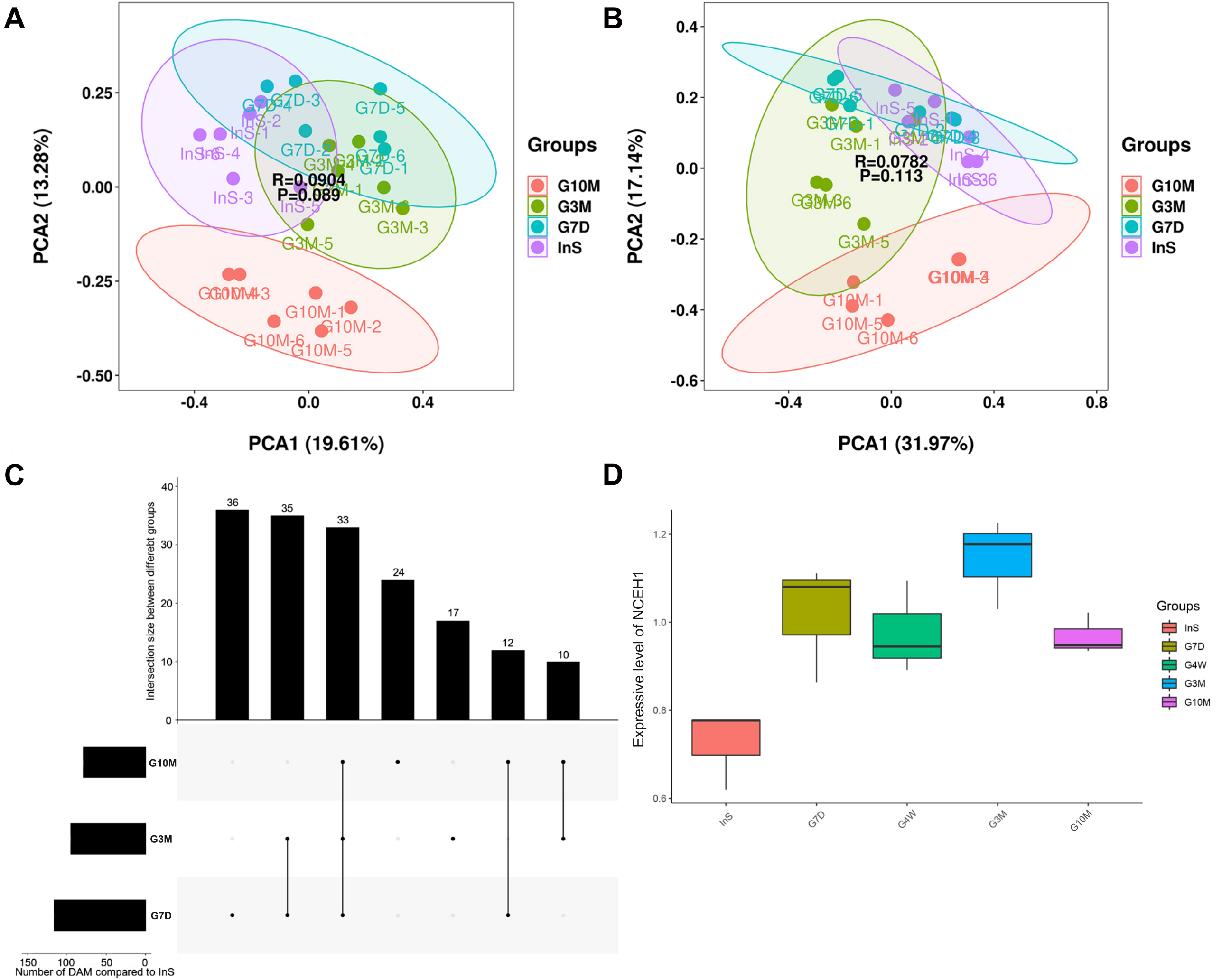

### Fig. S6

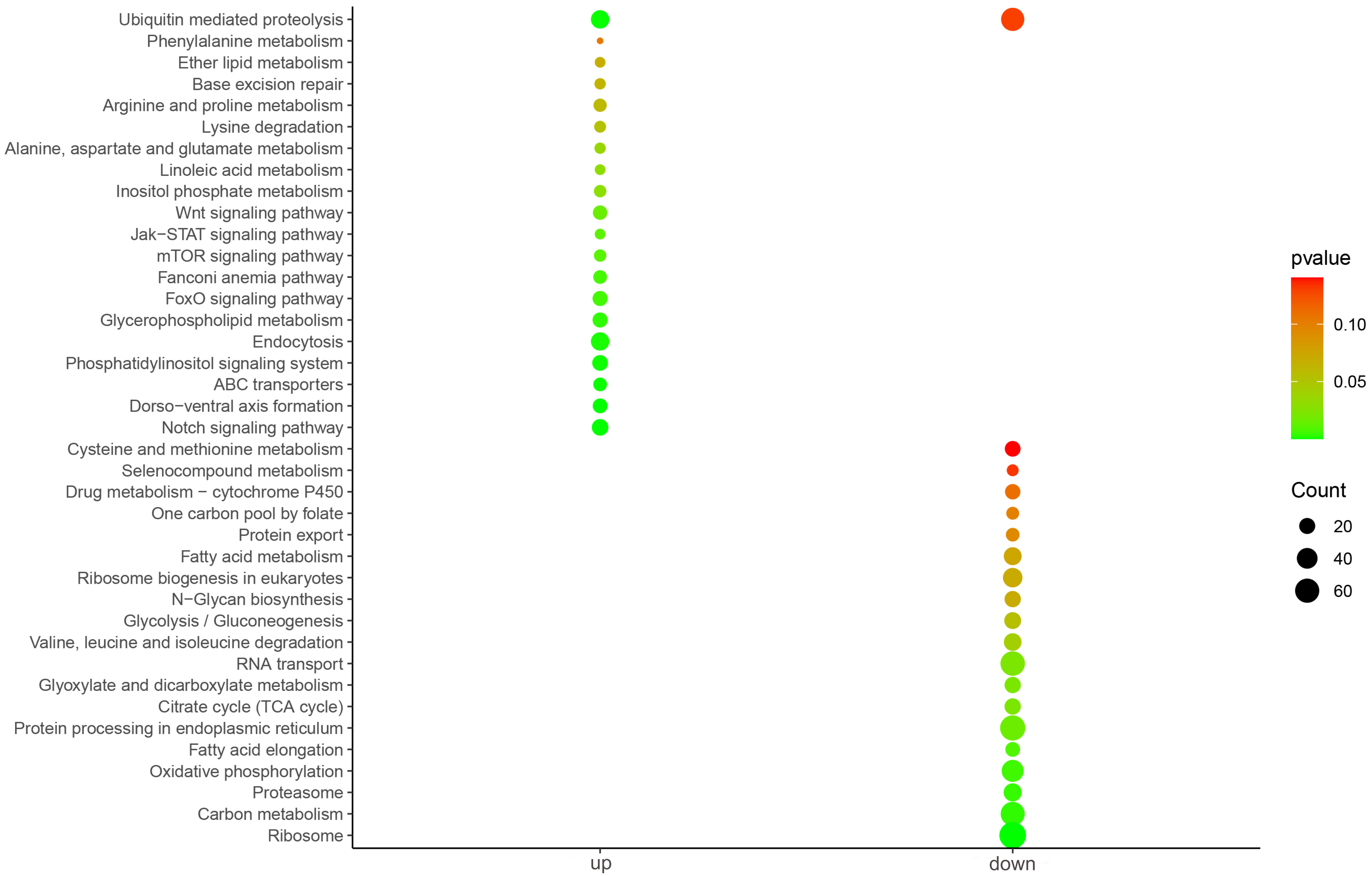

### Fig. S7

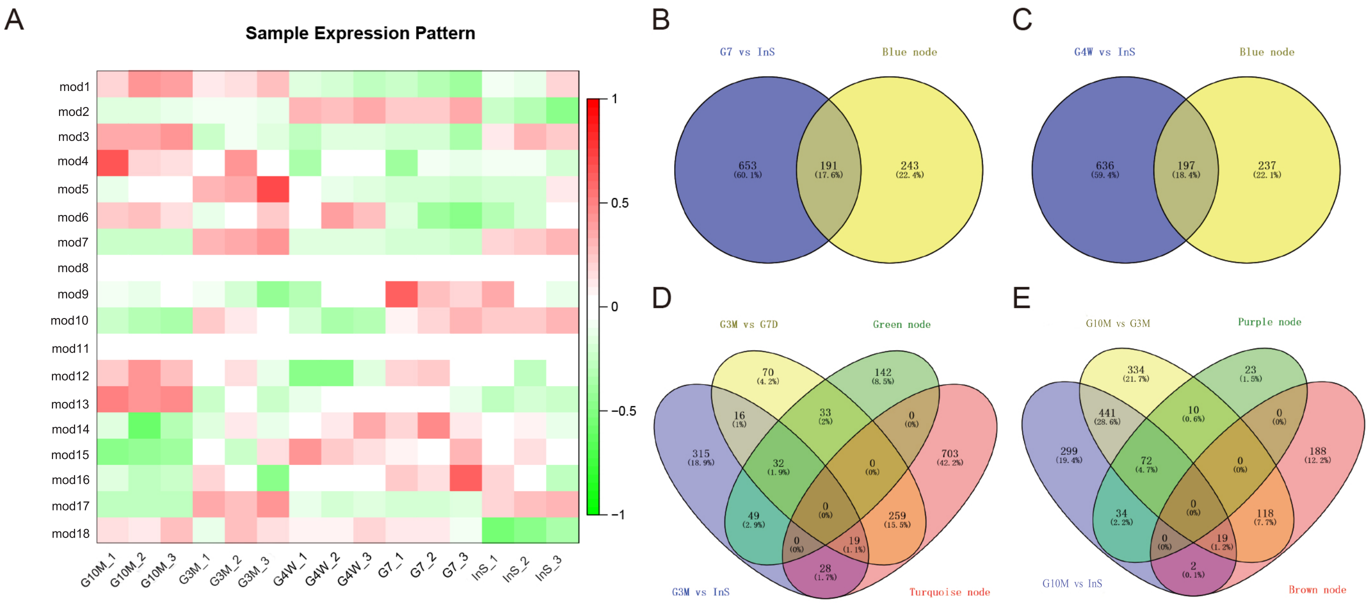

### Fig. S8

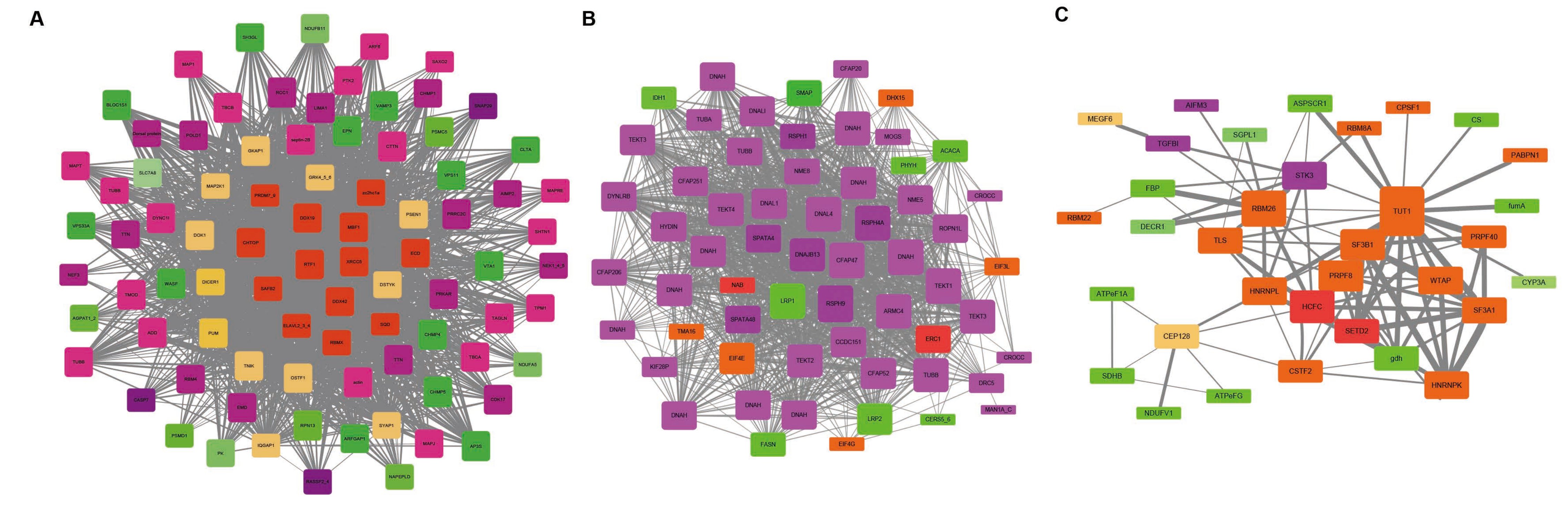
