## Supplementary text for "Phenotypic Plasticity of Symbiotic Organ Highlight Deep-sea Mussel as Model Species in Monitoring Exploitation of Deep-sea Methane Hydrate"

**Supplementary results**

***Statistics of the proteome and metabolome of gill tissue after symbiont depletion***

A total of 7821 host proteins were identified in the proteome, occupying 22.58% of total encoding genes in *G. platifrons*, and 878 endosymbiont proteins occupying 14.21% of their total encoding genes were identified from the proteome data of all groups (Table S2). A total of 844, 833, 459, and 867 differentially expressed proteins were identified in G7D, G4W, G3M and G1Y group in comparison with InS group, respectively (Table S3). Noticeably, we found the differential proteins in G7D and G4W group were highly overlapped while the G3M group was clustered more closely to InS group than other groups with the least number of differential proteins (Fig. S2CE). In comparison, more proteins were found differentially regulated in the G1Y group. The drastic alternations in the expression pattern of mussel proteins demonstrated the vigorous response of host against symbiont depletion and indicated that the host might adopt different strategies during the long-term symbiont decolonization. In addition, KEGG and GO enrichment analysis showed that biological processes such as transcription/translation, lysosome-mediated biodegradation, endocytosis, metabolism of fatty acids via peroxisome, monocarboxylic acid catabolic process and bacteria invasion of host cells were vigorously modulated in G7D and G4W group (Fig. S3AB, S4AB). In comparison, biological processes such as the transmembrane transport of metabolites, cilium assembly and movement and cell proliferation/differentiation began to be modulated in G3M group (Fig. S3C, S4C). As for the G1Y group, biological processes including transcription/translation, glucose metabolism, longevity regulating pathway, and amino acid metabolism were vigorously regulated (Fig. S3D, S4D). To further verify the metabolic response of deep-sea mussels, metabolomes of InS, G7D, G3M and G1Y group were also profiled (Table S4). PCA analysis showed that the InS group, G3M group and G1Y group were clustered distinctly with each other while G7D group interspersed between the InS and G3M group (Fig. S5AB). The differentially abundant metabolites identified by OPLS-DA test also showed distinct metabolic pattern between groups (Fig. S5C, Table S5), collectively suggesting that the mussel hosts were undertaking different responses to the long-term methane deprivation and the simultaneous symbiont decolonization.

***Meta-transcriptome analysis of mussels treated with antibiotics in situ***

To rule out the effect of the antibiotic treatment, we conducted meta-transcriptome analysis of gill tissue of six InS mussels and six *in situ* antibiotic treated mussels to assess their expression patterns. Consequently, a drastic decrease in the genome mapping rate of endosymbionts were observed, implying a global suppression of transcription process on endosymbionts. Besides, a total of 9197 host genes were found differentially expressed after the 7-day *in situ* antibiotic treatment (including 4655 up-regulated genes and 4542 down-regulated genes), while 3660 symbiont genes were differentially expressed, in which most of genes (3537 genes) were down-regulated (Supplementary Table S5). KEGG enrichment analysis of all the differentially expressed genes were subsequently conducted. In detail, host pathways involving signaling pathways, endocytosis and amino acid metabolism were significantly up-regulated, while glycometabolism and lipid metabolism were significantly down-regulated, which were inconsistent with the proteomic results of G7D or G4W group (Fig.S6, Supplementary Table S1). In addition, the up-regulation of lysosome-related pathways in G7D/G4W group was not observed in Bathymodioline mussels treated with antibiotics alone (Fig. S6, Table S1). Besides, glycolysis/gluconeogenesis, the citrate cycle (TCA cycle), and the fatty acid metabolism pathway were significantly down-regulated in mussels treated with antibiotics alone (Fig. S6, Table S1).

It was also noticeable that the relative abundance of symbionts remained comparable in control mussels (InS group) and in mussels treated with antibiotics alone (conducted *in situ* for 7 days, Fig. S2A) or acclimated under high pressure (2 MPa with methane, Fig. S2B), which confirmed the substantial influence of methane deprivation in decolonizing the deep-sea mussels.

**Supplementary Methods**

**Proteome analysis**

Gill samples employed for the proteome analysis were first grinded in liquid nitrogen and then treated with lysis buffer (8 M urea and 1% protease inhibitor cocktail) to release proteins. After sonication three times on ice using a high intensity ultrasonic processor (Scientz), cell debris was removed by a 10 min’s centrifugation at 12,000 g, 4 ℃ while the remaining supernatant was quantified with BCA protein assay kit. The obtained protein samples were then reduced with 5 mM dithiothreitol for 30 min at 56 ℃ and alkylated with 11 mM iodoacetamide for 15 min at room temperature in darkness. Afterward, TEAB solution (100 mM) was supplemented into the protein samples to dilute the urea to a finally concentration of less than 2M. Finally, the protein samples were digested by trypsin for two times (2% trypsin of total protein for the first time, overnight and 1% trypsin for the second time, 4 h) and subjected to the TMT labeling.

For the TMT/iTRAQ labeling assay, peptides obtained by trypsin digestion were first desalted by Strata X C18 SPE column (Phenomenex) and vacuum-dried. After dissolved in 0.5 M TEAB, the peptides were then incubated with TMT (in acetonitrile) for 2 h at room temperature according to the manufacturer’s protocol. After the labeling assay, the peptide mixtures were then pooled, desalted, dried by vacuum centrifugation and then fractionated (separated with a gradient of 8% to 32% acetonitrile, pH 9.0, 60 min) into 6 fractions by high pH reverse-phase HPLC using Thermo Betasil C18 column (5 μm particles, 10 mm ID, 250 mm length). After dried by vacuum centrifuging, the tryptic peptides were dissolved in 0.1% formic acid and separated using a home-made reversed-phase analytical column (15-cm length, 75 μm i.d.) by the EASY-nLC 1000 UPLC system. The peptides were finally subjected to NSI source followed by tandem mass spectrometry (MS/MS) in Q ExactiveTM Plus (Thermo) coupled online to the UPLC.

The resulting MS/MS data were processed using Maxquant search engine (v.1.5.2.8). Tandem mass spectra were searched against the *G. platifrons* protein database (generated from the updated genome information) concatenated with reverse decoy database. Trypsin/P was specified as cleavage enzyme allowing up to 4 missing cleavages. The mass tolerance for precursor ions was set as 20 ppm for the first search and 5 ppm in Main search, and the mass tolerance for fragment ions was set as 0.02 Da. Carbamidomethyl on Cysteine was specified as fixed modification and acetylation modification and oxidation on Met were specified as variable modifications. FDR was adjusted to < 1% and minimum score for modified peptides was set > 40.

GO annotation proteome was derived from the UniProt-GOA database (http://www.ebi.ac.uk/GOA/). Firstly, converting identified protein ID to UniProt ID and then mapping to GO IDs by protein ID. If some identified proteins were not annotated by UniProt-GOA database, the InterProScan soft would be used to annotated protein’s GO functional based on protein sequence alignment method. Then proteins were classified by Gene Ontology annotation based on three categories: biological process, cellular component and molecular function. KEGG database was used to annotate protein pathway. Firstly, using KEGG online service tools KAAS to annotated protein’s KEGG database description. Then mapping the annotation result on the KEGG pathway database using KEGG online service tools KEGG mapper. The GO / KEGG with a corrected p-value < 0.05 is considered significant.

**Metabolome analysis**

Gill tissues were first grinded in liquid nitrogen and then mixed thoroughly with 80% methanol to release all metabolites. After centrifuged at 12,000 g, 10 ℃ for 15 min, the supernatants were divided into four parts and subjected to LC-MS, GC-MS, QC of LC-MS, and QC of GC-MS correspondingly. All samples were first lyophilized for while the lyophilized supernatants for LC-MS were dissolved in 80 µL 25% acetonitrile aqueous solution before performing the LC-MS. For GC-MS, the lyophilized samples were first subjected to derivatization by oximation with methoxypyridine (50 µL, 20 mg/mL) for 1.5 h at 37 ℃. Samples were then silanized with 40 µL N-Methyl-N-(trimethylsilyl)trifluoroacetamide for 1 h at 37 ℃. After centrifuged at 12,000 g, 4 ℃ for 10 min, the derivative samples were then subjected to GC-MS.

For the LC-MS, samples were analyzed at both positive and negative ionization mode by Vanquish UPLC-Q-Exactive (Thermo Fisher Scientific, Rockford, IL, USA) to maximize metabolome coverage. For the positive ionization mode, metabolites were separated using Waters BEH C8 column (100 mm × 2.1 mm, 1.7 μm) with gradient formic acid solvent at a constant flow velocity of 0.35ml/min. The electrospray voltage applied was 3.5 kV and the m/z scan range was 70 to 1000 for full scan. Metabolites were detected in the Orbitrap at a resolution of 1.4×10^5^. For the negative ionization mode, Acquity UPLC HSS T3 column (100 mm × 2.1 mm, 1.8 μm) was used with gradient methanol solvent. The electrospray voltage applied was 3.0 kV and the m/z scan range was 80 to 1000 for full scan. A set of standard metabolites including Carnitine C2:0-d3, Carnitine C10:0-d3, Carnitine C16:0-d3, LPC 19:0, FFA C16:0-d3, FFA C18:0-d3, CA-d4, CDCA-d4, Phe-d5 and Trp-d5 were employed as internal control.

For the GC-MS, samples were analyzed by Agilent GC-MS 5977A with DB-5MS column (30m×0.25mm, 0.25μm). The temperature of inlet and ion sources was set at 320 ℃ and 230 ℃ correspondingly while the oven temperature program was set starting from 80 ℃ (1min) to 210 ℃ (with a ramp rate of 30℃/min) to 320 ℃ (with a ramp rate of 20 ℃/min and maintained for 4min). The m/z scan range was set from 33 to 600 for full scan.

**TEM imaging**

For TEM, gill samples fixed by paraformaldehyde-glutaraldehyde were treated with 1.0% osmium tetroxide (OsO_4_) for staining and then dehydrated with alcohol. After infiltration with acrylic resin, the samples were finally embedded and sectioned with a thickness of 70 nm using an ultramicrotome (EM UC7, Leica, Vienna, Austria). After staining by uranyl acetate and lead citrate, the gill cell ultrastructure was imaged with a transmission electron microscope (HT7700, Hitachi, Tokyo, Japan).

**Supplementary Figures**

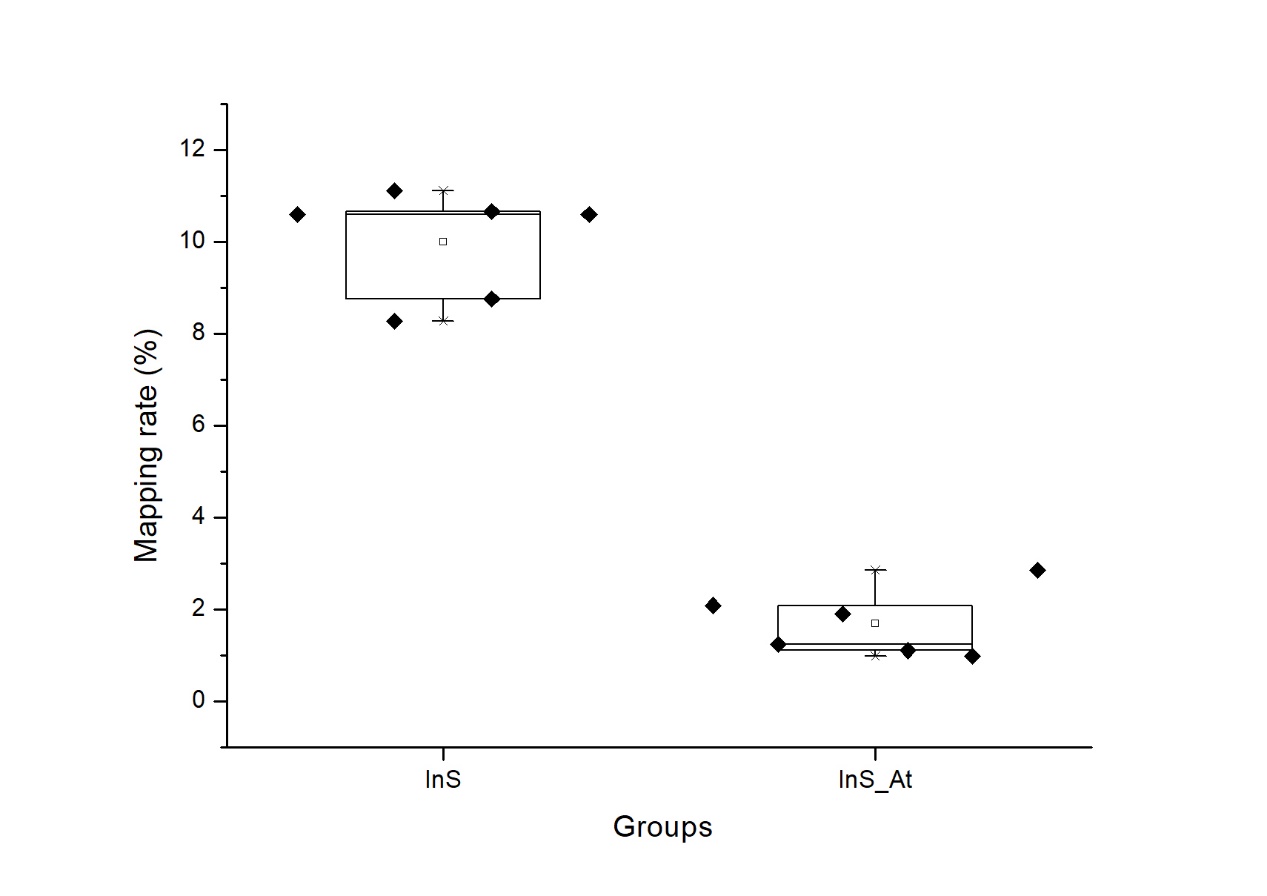

**Fig. S1. Gnome mapping rate of endosymbionts in mussels of InS and InS_At (*in situ* antibiotics treatment) group.**

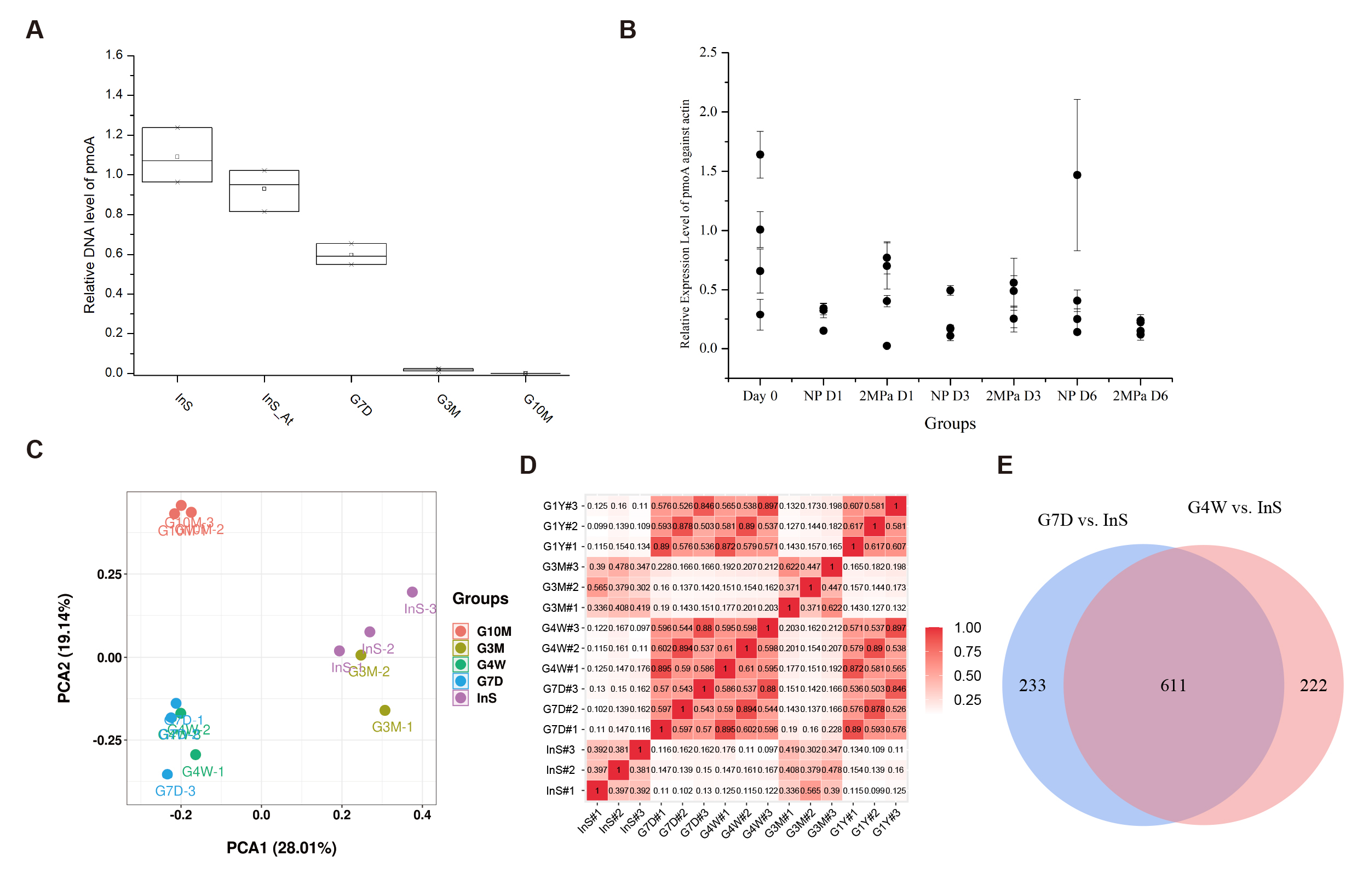

**Fig. S2. Overview of gill proteome under methane deprivation.**

**(A)** The relative abundance of the *pmoA* gene in the InS, InS_At (*in situ* antibiotic treatment), G7D, G3M and G10M groups.

**(B)** Principal component analysis (PCA) of proteome data in all groups.

**(C)** Pearson correlation analysis of the proteome in all groups (red indicates high correlation, white indicates low correlation).

**(D)** Venn diagram of differentially expressed proteins in the G7D and G4W groups compared to the InS group.

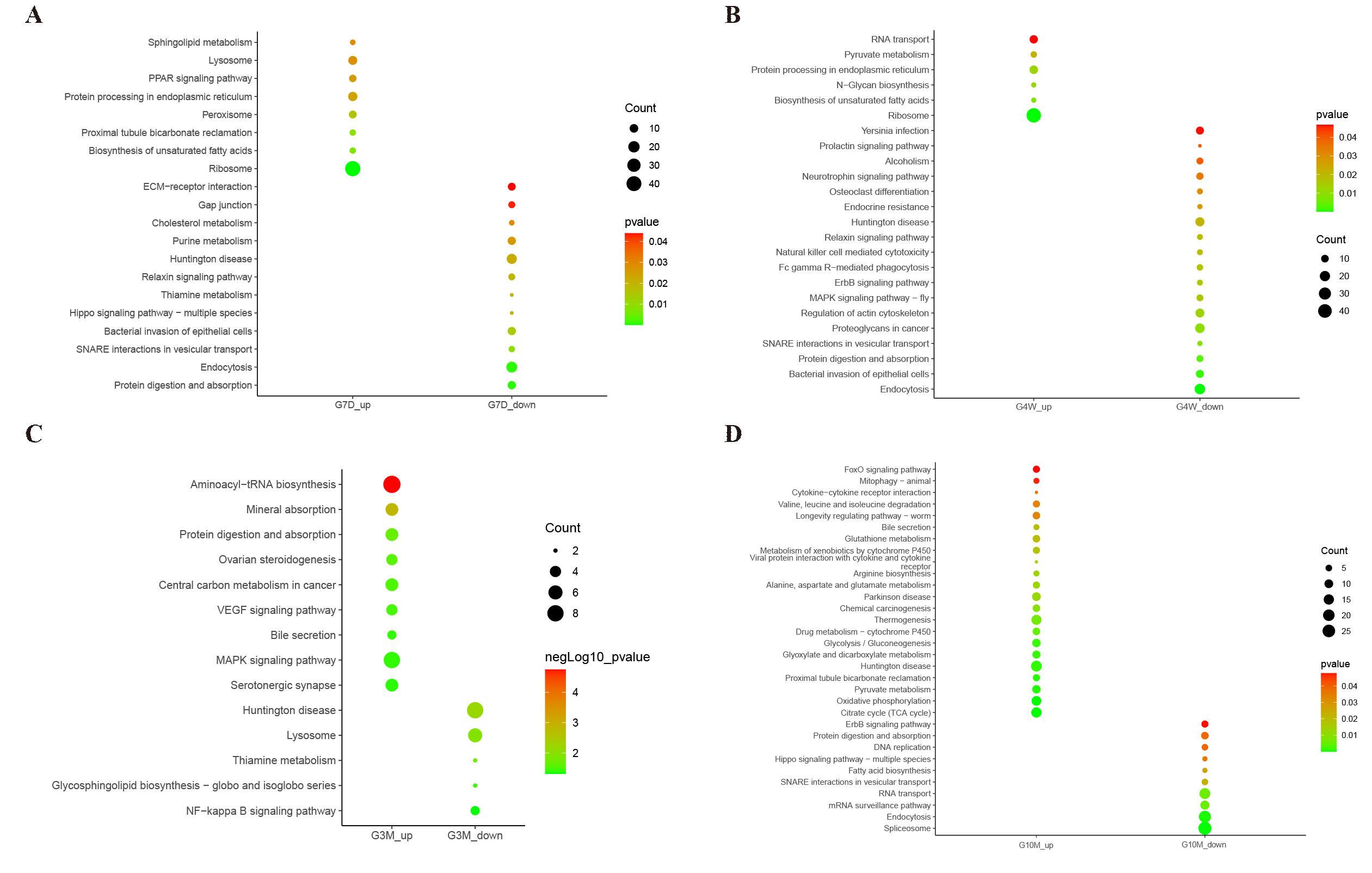

**Fig. S3.** **Kyoto Encyclopedia of Genes and Genomes (KEGG) enrichment analysis of differentially expressed proteins.**

The significantly enriched KEGG pathways of up-regulated and down-regulated differentially expressed proteins in the G7D group (A), G4W group (B), G3M group (C), and G10M group (D) compared to the InS group are illustrated.

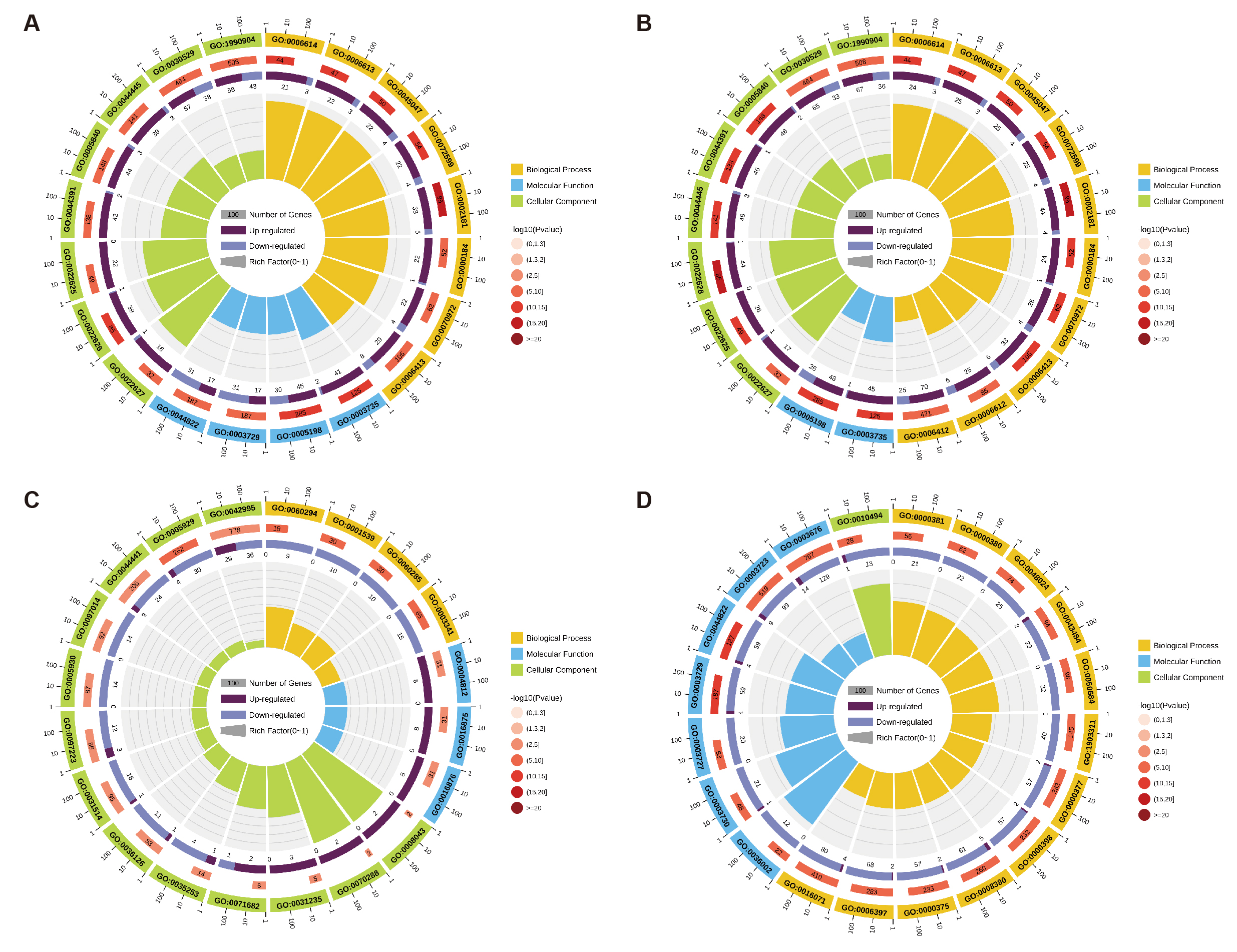

**Fig. S4. Gene Ontology (GO) enrichment analysis of differentially expressed proteins in the G7D, G4W, G3M, and G10M groups compared to the InS group.**

The top 20 most significant GO terms of the differentially expressed proteins of the G7D group (A), G4W group (B), G3M group (C), and G10M group (D) compared to the InS group are illustrated. The circles from outer to inner indicate the classification of GO terms (yellow indicates biological process, blue indicates molecular function, and green indicates cellular component), the number of background proteins in each classification (longer indicates more proteins), colored is used to indicate the *p*-value (red indicates greater significance), the numbers of up-regulated (intense purple) and down-regulated proteins (light purple), and the rich factor value of each classification (each compartment represents 0.1), respectively.
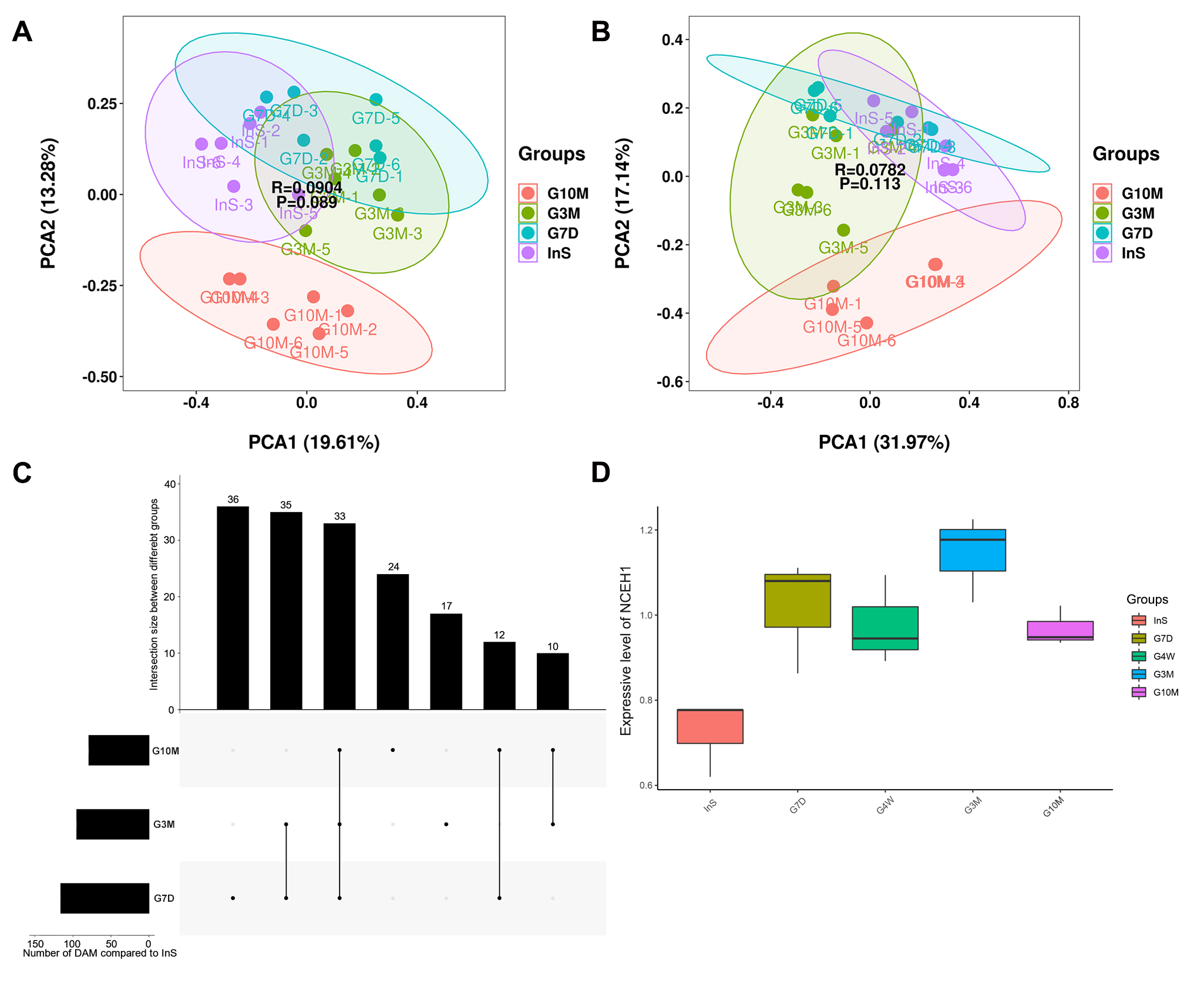
 **Fig. S5. Overview of metabolome analysis in the InS, G7D, G3M, and G10M groups.**

**(A)** Principal component analysis (PCA)of metabolites identified in the InS, G7D, G3M, and G10M groups by liquid chromatography–mass spectrometry (LC-MS).

**(B)** PCA analysis of metabolites identified in the InS, G7D, G3M, and G10M groups by gas chromatography–mass spectrometry (GC-MS).

**(C)** UpSet diagram of the differentially expressed proteins in the G7D, G3M, and G10M groups compared to the InS group. The horizontal bars show the total number of differentially abundant metabolites in each methane deprivation treatment group compared to the InS group. The vertical bars show the shared number of differentially abundant metabolites between the given groups. Connected dots refer to the groups involved in each intersection.

**(D)** Expression pattern of neutral cholesterol ester hydrolase 1 (NCEH1) during methane deprivation (data given as the mean ± SD).

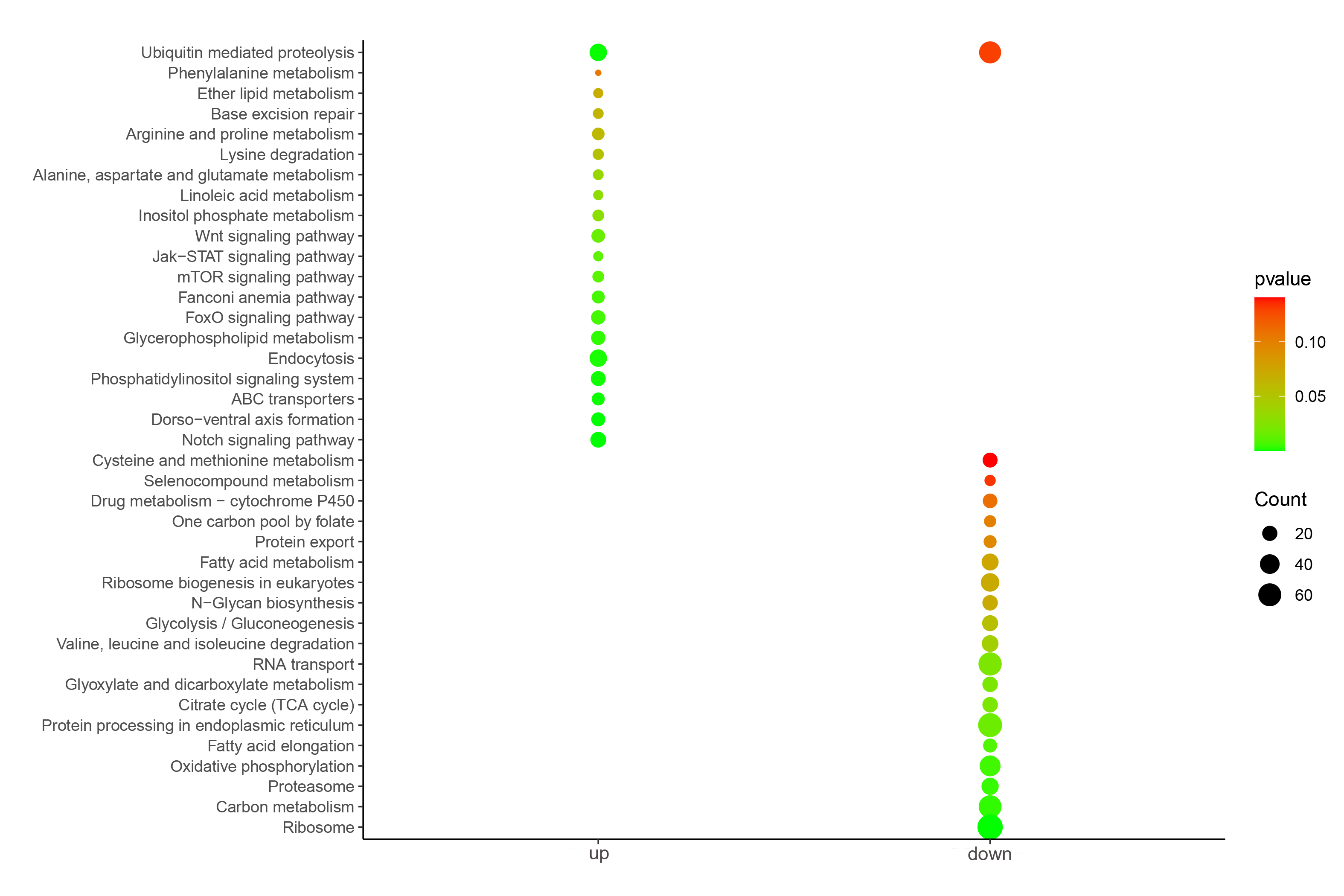

**Fig. S6. KEGG enrichment analysis of differentially expressed genes in *in situ* antibiotic treated mussels compared to InS mussels revealed by transcriptome.**

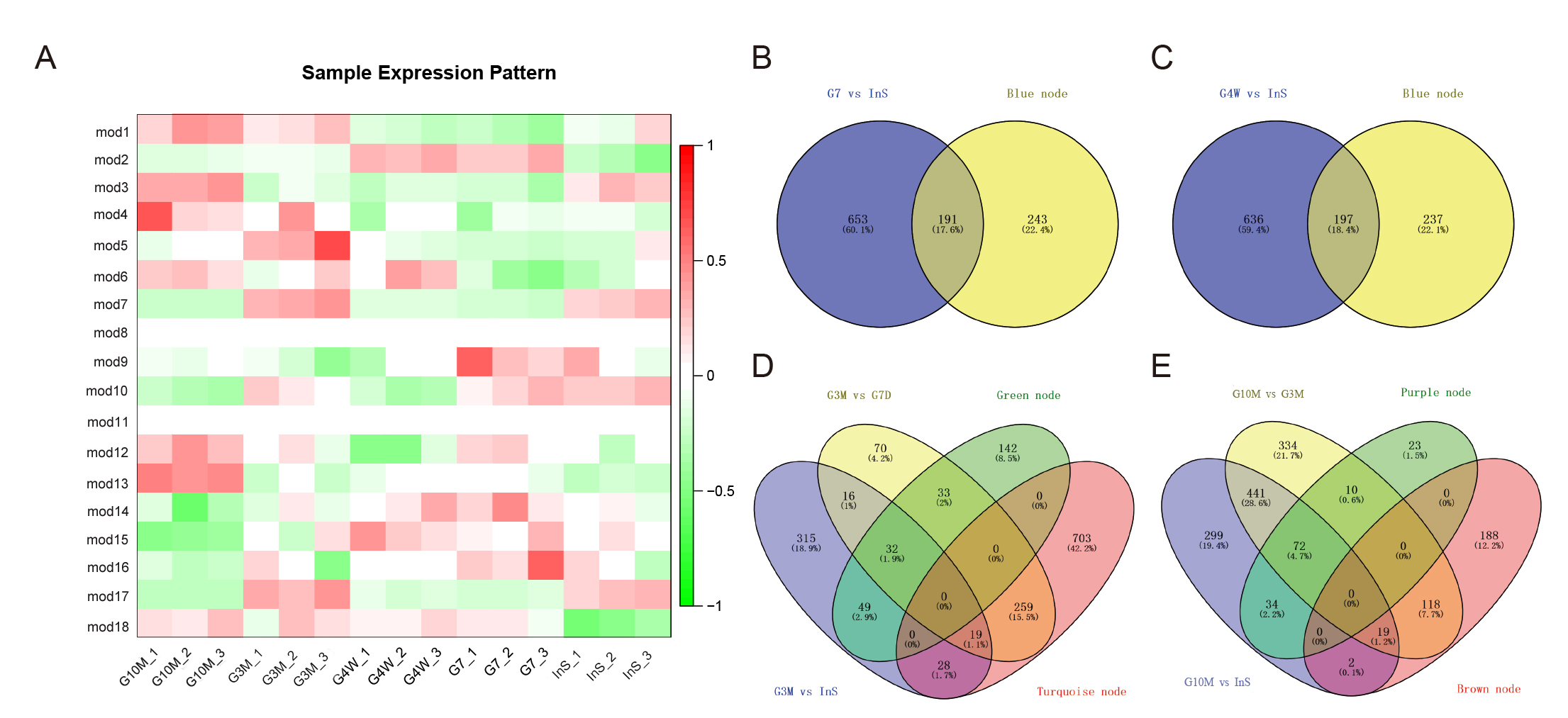

**Fig. S7. Sample expression pattern and venn diagrams of differentially expressed proteins identified in the G7D, G4W, G3M, and G10M groups and by weighted gene co-expression network analysis (WGCNA) analysis.**

**(A)** WGCNA Sample expression pattern revealed a total of 18 co-expression modules (mod1-18).

**(B)** Venn diagram of differentially expressed proteins in the G7D group and in the mod2.

**(C)** Venn diagram of differentially expressed proteins in the G4W group and in the mod2.

**(D)** Venn diagram of differentially expressed proteins in the mod5, mod17, and in the G3M group.

**(E)** Venn diagram of differentially expressed proteins in the mod13, mod3, and in the G10M group.

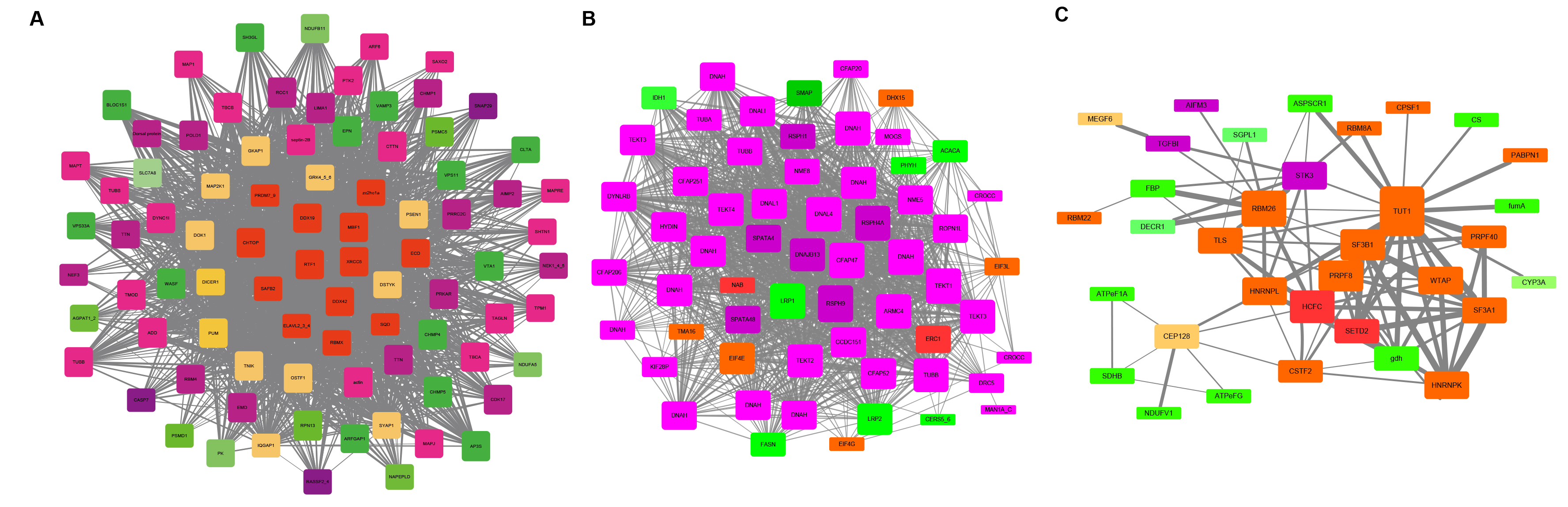

**Fig. S8. Weighted gene co-expression network analysis (WGCNA) of regulatory networks in the mod17, mod3, and mod13.**

The protein regulatory networks in the mod17 (A) and mod3 (B) in G3M group and mod13 (C) in G10M group constructed by WGCNA are illustrated.

**Supplementary Tables**

**Supplementary Table S1. List of identified proteins in the proteome of *G. platifrons* gills.**

**Supplementary Table S2. Differentially expressed proteins identified in different groups.**

**Supplementary Table S3. Identified metabolites in the metabolome of *G. platifrons* gills.**

**Supplementary Table S4. Differential metabolites identified in different groups.**

**Supplementary Table S5. Differential expressed genes in the meta-transcriptome analysis.**

**Supplementary abbreviations**

NRBF2: nuclear receptor binding factor 2; HSD17B4: (3R)-3-hydroxyacyl-CoA dehydrogenase; CYP39A1: 24-hydroxycholesterol 7alpha-hydroxylase; NCOA6: nuclear receptor coactivator 6; TCA: tricarboxylic acid cycle, ATP: adenosine triphosphate; CE: cholesterol ester; TG: triglyceride; PC: phosphatidylcholine; PE: phosphatidylethanolamine; SLC family: solute carrier family; NCEH1: neutral cholesterol ester hydrolase 1; CYP46A1: cholesterol 24-hydroxylase; NR1F4: nuclear receptor subfamily 1 group F member 4; NR1IN: nuclear receptor subfamily 1 group I; COL6A: collagen type VI alpha; ARS: aminoacyl-tRNA synthetase; EDF1: endothelial differentiation-related factor 1; CHTOP: chromatin target of PRMT1 protein; RTF1: RNA polymerase-associated protein RTF1 homolog; ECD: protein ecdysoneless homolog; ZNF260: zinc finger protein 260; EGFR: epidermal growth factor receptor; VEGF: vascular endothelial growth factor signaling pathway; MAPK: mitogen-activated protein kinase signaling pathway; pfkA: 6-phosphofructokinase 1; ALDO: fructose-bisphosphate aldolase; ENO: enolase; pdhB: pyruvate dehydrogenase E1 component beta subunit; MDH1: malate dehydrogenase; fumA: fumarate hydratase; SDHA/SDHB: succinate dehydrogenase (ubiquinone); LSC1/2: succinyl-CoA synthetase beta subunit; OGDH: 2-oxoglutarate dehydrogenase; IDH: isocitrate dehydrogenase; ACO: aconitate hydratase; CS: citrate synthase; GOT1: aspartate aminotransferase; ALT: alanine transaminase; PC: pyruvate carboxylase; OAT: ornithine--oxo-acid transaminase; PCD: 1-pyrroline-5-carboxylate dehydrogenase; GGT_5: gamma-glutamyltranspeptidase; GPT: glutathione S-transferase; GLUD1_2: glutamate dehydrogenase; NADH: nicotinamide adenine dinucleotide, reduced; NAD^+^: nicotinamide adenine dinucleotide, oxidized; FADH_2_: flavin adenosine dinucleotide, reduced; FAD: flavin adenosine dinucleotide, oxidized; TBC1D15: TBC1 domain family member 15; LC3: GABA (A) receptor-associated protein; OPTN: optineurin; ERC1: ELKS/Rab6-interacting/CAST family member 1; HCFC1: host cell factor 1; SPEN: protein split ends; TUT1: translation regulators speckle targeted PIP5K1A-regulated poly(A) polymerase.
